# Where single-cell foundation models help: a public-data benchmark of annotation, in silico perturbation and gene regulatory inference

**DOI:** 10.64898/2026.11.03.763968

**Authors:** Francesca Cupaioli, Ivan Merelli

## Abstract

Single-cell foundation models (FMs) are proposed as general tools for cell-type annotation, in silico perturbation and gene-regulatory-network inference. We benchmarked five FMs (scGPT, Geneformer V1 and V2, Nicheformer, scFoundation), four newer FMs, CellOracle, supervised perturbation models and task-matched simple baselines on public data, from Perturb-seq screens of cell lines and primary T cells to a brain organoid screen, the developing mouse cortex and lung cancer biopsies, under an analysis plan fixed before the new results. FMs helped most when labels were transferred across species: Nicheformer reached 0.98 accuracy from human to mouse lung, above every baseline (at most 0.94), and fine-tuned Geneformer V2 reached macro-F1 0.88 from mouse to human, against at most 0.74. In K562, zero-shot perturbations captured the direction of real responses (up to 76% of perturbations), Geneformer V2 tracked CRISPRa effect sizes with its overexpression protocol (ρ = 0.30), and its networks recovered tumour differential expression better than co-expression. Within a species, FM embeddings did not exceed principal components. In silico deletions followed target expression and, across seven screens with replicate-level ground truth, met the predefined criterion in none of 81 tests, although in K562 and RPE1 the measured effects replicated between libraries. In contrast, the gene vectors of seven of eight FMs, including three newer models, added to expression features, predicted which knockdowns have large effects in an independent genome-wide K562 screen (ρ between 0.34 and 0.44, against 0.26 with expression alone). We release the benchmark and its checks as the fmaudit package.

**Key Points:**

- Five single-cell foundation models, a GRN-based simulator, supervised perturbation models including STATE, and non-foundation baselines were compared on annotation, perturbation and GRN inference across cell lines, primary cells, genome-scale CRISPRi screens in primary T cells, a public organoid CRISPR screen, an in vivo mouse cortex screen and human tumour biopsies, all from public data.
- Foundation models were most useful for transferring cell-type labels across species. From human to mouse Nicheformer exceeded every baseline, and fine-tuned Geneformer V2 did so from mouse to human. Within a species principal components matched or exceeded every model, also after fine-tuning.
- Zero-shot in silico perturbations captured the direction of real K562 responses for several models, and with its native overexpression protocol Geneformer V2 also tracked CRISPRa effect sizes. In an organoid screen and in the mouse cortex, predicted knockout effects followed target expression and were no larger than those of expression-matched control genes. In genome-scale screens of primary CD4 T cells, K562 and RPE1, predicted effect sizes did not exceed target expression or the number of perturbed cells, although the measured effects replicated well in the two cell lines, and their shifts pointed towards the real response of the deleted gene only faintly (2 of 20 tests). The gene vectors that FMs learn, however (seven of eight models tested, including three newer ones), predicted which knockdowns have large effects beyond expression, as confirmed on an independent genome-wide K562 screen, and remained informative over STRING degree in a robustness analysis.
- With patient-level ground truth, Geneformer V2 networks recovered tumour differential expression better than co-expression. Against measured perturbation effects in neural tissue, no model exceeded naive predictors such as the number of perturbed cells, and the measured effects themselves replicated only partly between libraries.
- Benchmarks of single-cell foundation models should include task-matched simple baselines, replicate-level ground truth, permutation nulls within cell type and an expression-matched control for in silico knockouts. We release the baselines, checks and specificity test as the fmaudit package, with the permutation null as an analysis script.

## Introduction

Transformer models pre-trained on tens of millions of single-cell transcriptomes, such as scGPT [1], Geneformer [2], Nicheformer [3] and scFoundation [4], are presented as foundation models (FMs) that can be applied, with or without fine-tuning, to many downstream tasks, most often cell-type annotation, prediction of the transcriptional response to genetic perturbation, and inference of gene regulatory networks (GRNs). If reliable, FMs would enable in silico experiments in systems that are expensive to modify genetically, such as organoids and tissues.

Independent evaluations have tempered these expectations. For clustering and data integration, zero-shot FM embeddings often perform no better than highly variable genes or principal components [5,6]. For perturbation prediction, deep-learning models, FMs included, do not outperform deliberately simple linear or mean baselines [7,8], and neither zero-shot FM embeddings [9] nor expression-forecasting methods [10] improve on them. These studies mostly evaluate one task in cell lines, where large Perturb-seq screens provide ground truth [11,12,13]. It remains unclear whether their conclusions extend to tissue-like systems and to the disease models in which in silico perturbation would be most useful, and whether a model that performs well on one task or system can be trusted on another.

Here we evaluate the same methods on three tasks (annotation, in silico perturbation and GRN inference) at three levels of biological complexity, using only public data: (i) cell lines and primary cells, including Perturb-seq screens of K562 and RPE1 [13] and a genome-scale CRISPRi screen of primary human CD4 T cells [14], (ii) neural tissue, with the CHOOSE mosaic CRISPR screen of 36 genes in brain organoids [15] and an in vivo Perturb-seq screen of 35 autism risk genes in the developing mouse cortex [16], together with CRISPRi and CRISPRa screens of iPSC-derived neurons [17], and (iii) biopsies of EGFR-mutant lung cancer. Perturbation screens in tissue provide the ground truth that in silico perturbation needs, if the measured effect of a gene can be separated from differences between cell types and libraries. We therefore fixed the analysis plan before computing the new results, scored every prediction against naive predictors and permutation nulls within cell type, and estimated how well the measured effects replicate between libraries.

Errors in data scaling or model training are easy to introduce and rarely visible in the results. We therefore checked every analysis step: input-scale checks, verification of weight changes after fine-tuning, train/test overlap checks, target-detection checks and fixed model inputs for in silico knockouts. These checks changed several conclusions, and we release them in the fmaudit package.

## Methods

### Benchmark design

The benchmark is a grid of tasks (annotation, perturbation, GRN) by datasets at three levels of biological complexity (Figure 1). Methods were scGPT [1], Geneformer V1-10M [2] and V2-316M, Nicheformer [3] and scFoundation [4], used zero-shot with the input processing of their own tutorials and, where applicable, after fine-tuning. We also ran CellOracle [18] with a promoter base GRN, optionally augmented with STRING v12 interactions [19], the supervised perturbation models GEARS [20], CPA [21], scGPT fine-tuned for perturbation and STATE [22], and non-FM baselines chosen per task, including gene-independent predictors such as the number of cells per cell type. All methods in a grid row were run on the same cells, split, gene universe and metric. Full protocols are in the Supplementary Methods.

**Figure 1.**
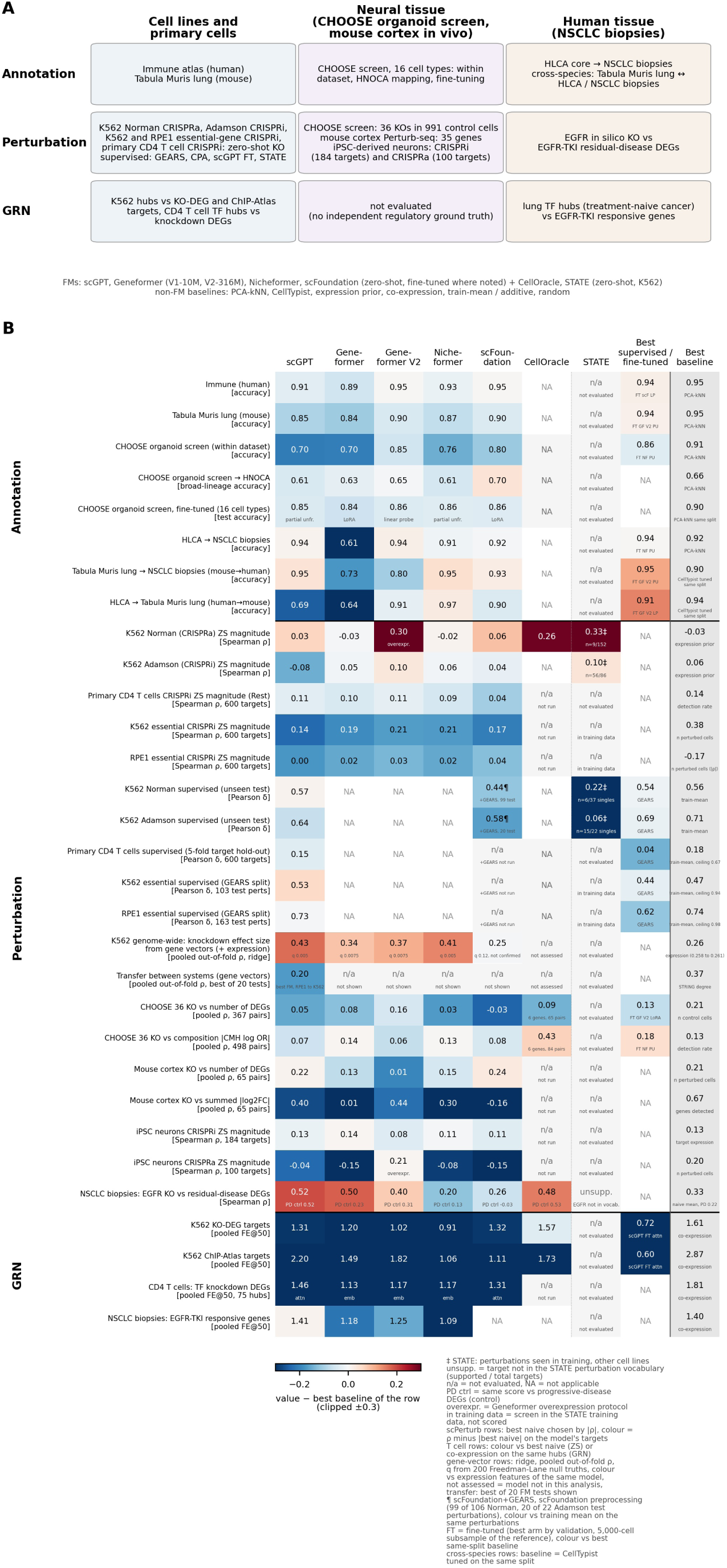
Benchmark overview. (A) Tasks, datasets and methods at three levels. The first level covers cell lines and primary cells: K562 Norman CRISPRa, Adamson CRISPRi, K562 and RPE1 essential-gene CRISPRi and the primary CD4 T cell CRISPRi screen. The second covers neural tissue: the CHOOSE organoid screen, the in vivo mouse cortex screen and the iPSC-derived neuron CRISPRi and CRISPRa screens. The third covers human tissue (NSCLC biopsies). (B) Scorecard of all task by dataset rows. Colour shows each method’s value minus the best non-FM baseline of the row, clipped at plus or minus 0.3, and text shows the value. Rows use different metrics, so colours compare methods within a row only. For the organoid screen and mouse cortex knockouts, values are pooled Spearman ρ over gene by cell-type (or class) pairs of the zero-shot predictions. The truths are the measured number of DEGs, the composition change (organoid screen) and the summed |log2FC| (mouse cortex, GFP reference). The baseline is the best naive predictor on the same pairs. The fine-tuned column of these rows gives the best of the eleven fine-tuned backbones. The CellOracle values of the organoid screen cover the six genes it can simulate and are coloured against the best naive predictor on the same pairs. For the primary CD4 T cell, K562 and RPE1 essential-gene and iPSC neuron CRISPRi and CRISPRa screens, values are Spearman ρ between the zero-shot deletion effect and the real number of DE genes over 600, 600, 600, 184 and 100 targets (T cells, resting cells against the resting truth). The baseline is the best naive predictor on all targets, and each model is coloured against the best naive predictor recomputed on its own targets. For the scPerturb screens the best naive predictor is chosen, and subtracted, by its absolute ρ, as pre-registered. The gene-vector rows report ridge regression on FM gene vectors, scored as pooled out-of-fold Spearman ρ with p and q from 200 Freedman-Lane null truths. In K562 genome-wide, the knockdown effect size predicted from gene vectors plus expression features reached ρ 0.43 for scGPT, 0.41 for Nicheformer, 0.37 for Geneformer V2 and 0.34 for Geneformer V1 (q at most 0.0075) and 0.25 for scFoundation (q 0.12, not confirmed), against 0.26 for the expression features alone (0.258 to 0.261 across the analyses). Each model is coloured against its own expression-only baseline (Supplementary Figure S9). In the transfer between systems, no FM exceeded the baseline in any of the 20 tests. The best FM test is shown (scGPT, RPE1 to K562, ρ 0.20), against the best baseline, STRING degree (ρ 0.37). In the supervised K562 Norman and Adamson rows the scFoundation column is scFoundation with GEARS (¶ 99 of 106 Norman and 20 of 22 Adamson test perturbations, against the training mean on the same perturbations). In the supervised rows of the genome-scale screens the scGPT column is scGPT with GEARS, the best supervised column is GEARS and the baseline is the training mean, with the split-half ceiling in the sub-label (Pearson δ on the top 20 DE genes, five-fold target hold-out over 600 targets for T cells, GEARS split with 103 and 163 test perturbations for K562 and RPE1). The T cell GRN row gives the pooled expression-matched FE@50 of up to 75 transcription-factor hubs against the authors’ DE genes. Each FM value is the best of its embedding and attention networks (named in the sub-label) and is coloured against signed Pearson co-expression on the same hubs. The Geneformer V2 values of the K562 Norman and neuron CRISPRa rows use Geneformer’s overexpression protocol. In the annotation rows the fine-tuned column gives the best test accuracy over the five FMs fine-tuned as classifiers, each arm (linear probe, partial unfreezing or LoRA) chosen on validation on a 5,000-cell subsample of each reference, against the best baseline trained on the same cells. The two cross-species rows use CellTypist tuned on the same cells as baseline. The organoid screen fine-tuned row shows each model’s arm selected on validation against PCA-kNN trained on the same cells. PD ctrl is the same score against the progression control. ‡ STATE evaluated on perturbations seen in training in other cell lines. In training data means that STATE was not scored because the Replogle essential-gene screens are in its training data. Unsupp. means target not in the model vocabulary (supported/total targets), n/a not evaluated or not run, not assessed an FM not included in the gene-vector analysis, not shown an FM transfer test below the best one shown, and NA not applicable. Geneformer V1 and V2 are shown as separate columns.

### Datasets

Level 1, cell lines and primary cells: the Immune Human atlas (33,506 cells, 16 types) [23], Tabula Muris Senis lung (mouse) [24], and the Norman CRISPRa (91,205 cells) [11] and Adamson CRISPRi (68,603 cells, 87 conditions) [12] K562 Perturb-seq screens in GEARS processing, the CRISPRi essential-gene Perturb-seq screens of K562 and RPE1 [13] from the scPerturb collection [25], and a genome-scale CRISPRi Perturb-seq screen of primary human CD4 T cells from four donors, at rest and after 8 and 48 hours of stimulation [14]. Level 2, neural tissue: the CHOOSE mosaic organoid screen [15] (28,344 cells, 36 perturbed genes, 991 non-targeting control cells, 16 cell types annotated by the authors), in which perturbed and control cells share the same organoids, with HNOCA [26] as annotation reference, the CRISPRi and CRISPRa screens of iPSC-derived neurons [17] from the scPerturb collection, and the in vivo Perturb-seq screen of 35 autism and neurodevelopmental risk genes in the developing mouse cortex [16] (GEO GSE157977, perturbations delivered in utero at E12.5, cells profiled at P7), with the authors’ cell classes and perturbation assignments from the Single Cell Portal [27] (17,889 analysis cells, 17 libraries, five cell classes, 374 GFP control cells). Datasets and their processing are listed in Supplementary Table S1, and the design of the mouse cortex analysis in Supplementary Table S2.

Level 3, human tissue: EGFR-mutant non-small-cell lung cancer biopsies [28], with the HLCA core [29] as human lung reference.

### Annotation

scGPT used the continual-pretrained whole-human checkpoint and, for the organoid screen, also the original whole-human and brain checkpoints. Zero-shot cell embeddings were used for k-nearest-neighbour label transfer (k = 30 and cosine distance within a dataset, k = 20 and Euclidean distance with a confidence override across datasets). Cross-species transfer used one-to-one orthologs in both directions and a coarse crosswalk of shared classes, and was repeated with cosine kNN (k = 30, no override) as a classifier control. Cells left unassigned by the override were not scored (fractions and accuracy over all cells in Supplementary Table S3). Baselines were PCA on 2,000 highly variable genes with the same classifier, z-scored highly variable genes and CellTypist [30] trained on the reference. Organoid screen cells were annotated within the dataset (stratified 50/50 split) and, after mapping onto HNOCA, scored as broad-lineage concordance with the authors’ annotation.

### In silico perturbation

Zero-shot knockouts followed the embed, delete and re-embed protocol: the target gene was removed from each cell’s input, and the effect was the cosine distance between original and perturbed embeddings, averaged per cell type. For scGPT the sampled input genes and tie-breaks were fixed per cell, so that the two inputs differed only by the target. A deletion changes the input only in cells that express the target, so target detection was checked per cell type before simulation. In K562, each single-gene perturbation in a model’s vocabulary was applied to 500 control cells, and Geneformer was also run with its own protocol, which models CRISPRa by moving the target to the first rank of the input. In the organoid screen the 36 genes were deleted in all 991 non-targeting cells. In the mouse cortex the genes were deleted in the 374 GFP control cells from raw counts, through one-to-one human orthologs for the models trained on human data (32 of the 35 genes have an ortholog in our table) and with the mouse species token for Nicheformer. Supervised models were scored by Pearson δ on unseen GEARS test perturbations against training-mean and additive baselines [7]. GEARS, scGPT fine-tuned and GEARS with frozen scFoundation embeddings, the protocol of [4], were also trained on CHOOSE with five-fold gene hold-out, and Geneformer with its own perturbation-classifier protocol.

Because cells of one library are not independent replicates [31], measured effects in tissue were computed on pseudobulk counts per library and cell type with edgeR quasi-likelihood [32]. In the organoid screen the ground truth per gene and cell type was the number of DEGs at FDR < 0.05 and the absolute library-stratified Cochran-Mantel-Haenszel log odds ratio of composition (Supplementary Table S4). In the mouse cortex perturbed cells were compared with GFP control cells of the same class and library (paired design, at least 3 cells of each group per library and at least 20 perturbed cells over at least 2 libraries), giving 72 gene by class pairs, 65 of them with predictions, and, as a secondary reference, with all other perturbed cells of the same class and library (152 pairs, Supplementary Table S5). The measured effect was the number of DEGs at FDR < 0.05 and the summed absolute log2 fold change. The analysis plan, fixed before these results, called a prediction positive if its Spearman ρ over gene by cell-type pairs was positive with Benjamini-Hochberg q < 0.05 within its family, exceeded the best naive predictor by at least 0.10 with a bootstrap interval of the difference above 0, and exceeded a null in which perturbed genes were permuted within expression decile or within cell type (1,000 permutations). Naive predictors were the target’s mean expression and detection rate, the number of perturbed and control cells, and library size and genes detected per cell type. The replication ceiling was the agreement of the measured effects between two halves of the libraries.

Genome-scale screens. In the primary CD4 T cell screen [14], 600 targets were drawn by decile of target expression (seed 0) among the 5,217 targets with significant on-target knockdown and no off-target flag in all three culture conditions and scored in at least three donors, and deleted in 200 resting non-targeting cells (50 per donor). The measured effect was the authors’ number of DEGs (DESeq2, 10% FDR) at rest and after 8 and 48 hours of stimulation, the stimulated conditions being scored with the resting predictions, and the replication ceiling was the agreement between donors of per-donor edgeR effects. In the scPerturb screens [25], 600 targets shared by K562 and RPE1 were drawn by K562 expression decile, and all targets were used in the neuron screens. The measured effect was the number of genes with Wilcoxon p < 0.01 among the 3,000 most variable genes, perturbed against library-matched control cells, from raw counts, and the replication ceiling was the agreement between two halves of the libraries. Predictions used 500 control cells (all 437 and 434 in the neuron screens, and 100 for Geneformer V2 and scFoundation in K562, RPE1 and neuron CRISPRa because of their cost). The decision rule, the naive predictors (target expression, detection rate, number of perturbed cells) and the permutation within expression decile were those of the tissue screens, with Benjamini-Hochberg correction over the 15 T cell and the 20 cell-line tests. GEARS and scGPT with GEARS were trained on resting T cells with five-fold target hold-out and on the GEARS-processed essential-gene screens with the GEARS split (412 and 651 perturbations whose target is among the 5,000 genes of the files). STATE was not scored on these screens because they are in its training data.

Gene representations. The context-free gene vector of each FM (the input embedding of the gene, without positional embedding) was used as a feature of a ridge regression of the log number of DEGs of each target, alone or added to expression features (mean log expression and detection rate of the target in control cells, log number of perturbed cells) or to the STRING v12 degree of the target (combined score at least 700). Principal components (at most 50), standardisation and the penalty were fitted on training targets only. Transfer between systems used fixed halves of the shared targets, and analyses within a system used 5-fold (10-fold for the genome-wide screen) target cross-validation with pooled out-of-fold Spearman correlation. A code test on random truths showed that this procedure can produce gains of up to 0.05 by itself, so the gain over expression was tested against 200 null truths with permuted residuals of a least-squares fit on the expression features (Freedman and Lane), an amendment made before any score of the confirmatory screen [33]. The confirmatory screen was the genome-wide CRISPRi Perturb-seq of K562 [13] (7,568 targets with at least 25 perturbed cells, 267 libraries), with effects computed as for the essential-gene screens. Zero-shot deletion effects were also re-read without rerunning any model, as the effect per control cell expressing the target and as additional features of the same ridge model. The confirmatory scoring was repeated for Geneformer V1 and V2 and scFoundation with only the gene vectors changed (BH over these three tests). Direction specificity and the newer models are described in the Supplementary Methods. Partial knockdown was simulated by scaling the target count in the control cells by its residual expression measured in each screen.

For the lung-cancer biopsies [28] (23,261 Smart-seq2 cells), the ground truth was the number of DEGs per cell type for residual disease versus treatment-naive biopsies, per cell and, as a patient-aware control, by pseudobulk edgeR, with progression versus treatment-naive as specificity control. Library size and genes detected per cell type served as complexity baselines.

### Gene regulatory networks

For each FM, gene-gene networks were derived from contextual gene-embedding similarity and from attention averaged over layers and heads, and compared with CellOracle, Pearson co-expression, an expression prior and random ranking. The top-50 neighbours of each hub gene were scored against knockdown targets (Adamson) and ChIP-seq targets (ChIP-Atlas [34]) in K562, genes responding to EGFR inhibition in treatment-naive cancer cells of the lung-cancer biopsies, as fold enrichment over an expression-matched expectation (FE@50, 1 = chance), pooled over hubs. The same networks were derived from fine-tuned backbones, including each FM adapted to K562 with the hub perturbations held out.

### Fine-tuning

scGPT was fine-tuned end-to-end for perturbation on Norman or Adamson. Each FM was fine-tuned as a cell-type classifier on every annotation reference, including the organoid screen, with a linear probe, partial unfreezing and LoRA (scGPT also with the full fine-tuning of its tutorial), on 4,000 training and 1,000 validation cells, with the configuration selected on validation and tested once, against logistic regression on principal components (PCA-LR) or on z-scored variable genes, PCA-kNN and CellTypist tuned on the same cells. Knockouts were recomputed with each fine-tuned backbone, and tensor-wise weight checks confirmed which parameters changed. Networks were also derived from each FM adapted to K562 with the hub perturbations held out. Full settings are in the Supplementary Methods.

### Expression-matched specificity test

To test whether a knockout carries information about the deleted gene beyond the loss of expressed input, the embedding shift after deleting target g in a cell was compared with the shifts after deleting, one at a time, 20 control genes matched to g in that cell for expression and across the dataset for detection rate. The specific effect is S = observed − mean(control) and z = S / sd(control). A gene×cell-type pair with z > 2 has a specific excess shift. In the organoid screen the test was run on the 991 control cells for every zero-shot and fine-tuned backbone (Supplementary Table S6).

### Statistics and computation

Correlations are Spearman’s ρ with two-sided p-values unless stated otherwise. Multiple testing was controlled with the Benjamini-Hochberg procedure [35] within pre-specified families (K562, organoid screen annotation, organoid screen knockouts, mouse cortex, T cells, cell-line screens), and the number of tests is reported in the Supplementary Tables (for the organoid screen in Supplementary Table S7). The analyses of the organoid screen, the mouse cortex and the genome-scale screens followed an analysis plan written before their results (available in the repository), and every analysis run is reported. All computation ran on SLURM nodes (NVIDIA A100 and L40 GPUs) at CNR-ITB.

## Results

### A task-by-system map with baselines

Figure 1 summarises all task×dataset rows and method columns, each with a non-FM baseline computed on the same split and metric. In most rows the best FM was close to the baseline or below it. The exception was cross-species annotation.

### Annotation: foundation models help mainly across species

On primary immune cells, zero-shot kNN transfer from FM embeddings reached accuracies of 0.887 (Geneformer), 0.912 (scGPT), 0.926 (Nicheformer) and 0.949 (scFoundation). PCA with the same classifier reached 0.945 and CellTypist 0.921 (Figure 2A, Supplementary Table S8). On Tabula Muris lung, PCA (0.945, macro-F1 0.844) exceeded all FMs (0.839 to 0.902, macro-F1 0.635 to 0.765, Figure 2C, Supplementary Figure S1, Supplementary Table S9). Mapping the lung-cancer biopsies onto the HLCA, FMs (0.91 to 0.94, except Geneformer V1 at 0.605) were close to PCA (0.924) and CellTypist (0.913, Supplementary Table S10). From human to mouse (Supplementary Figure S2, Supplementary Tables S11 to S15), Nicheformer (0.968), Geneformer V2 (0.914) and scFoundation (0.898) exceeded CellTypist with default settings (0.684) and cosine-kNN baselines (0.863), but CellTypist with tuned regularisation reached 0.941 (macro-F1 0.883), so only Nicheformer remained above every baseline. From mouse to human, no FM was separable from cosine-kNN PCA (0.945). Only Nicheformer was pre-trained on mouse cells (Supplementary Table S16). Fine-tuned classifiers raised macro-F1 over zero-shot kNN within Tabula Muris lung for all five models (by 0.09 to 0.20), but only Geneformer V2 exceeded the best baseline trained on the same cells (0.892 versus 0.868). Across species fine-tuning lowered human-to-mouse macro-F1 below zero-shot for every model, whereas Geneformer V2 improved mouse-to-human transfer from 0.789 to 0.879, above every same-split baseline (at most 0.736, Supplementary Figure S3, Supplementary Table S17). With the full reference, zero-shot Nicheformer reached 0.894 and cosine-kNN PCA 0.862 (Supplementary Table S3).

**Figure 2.**
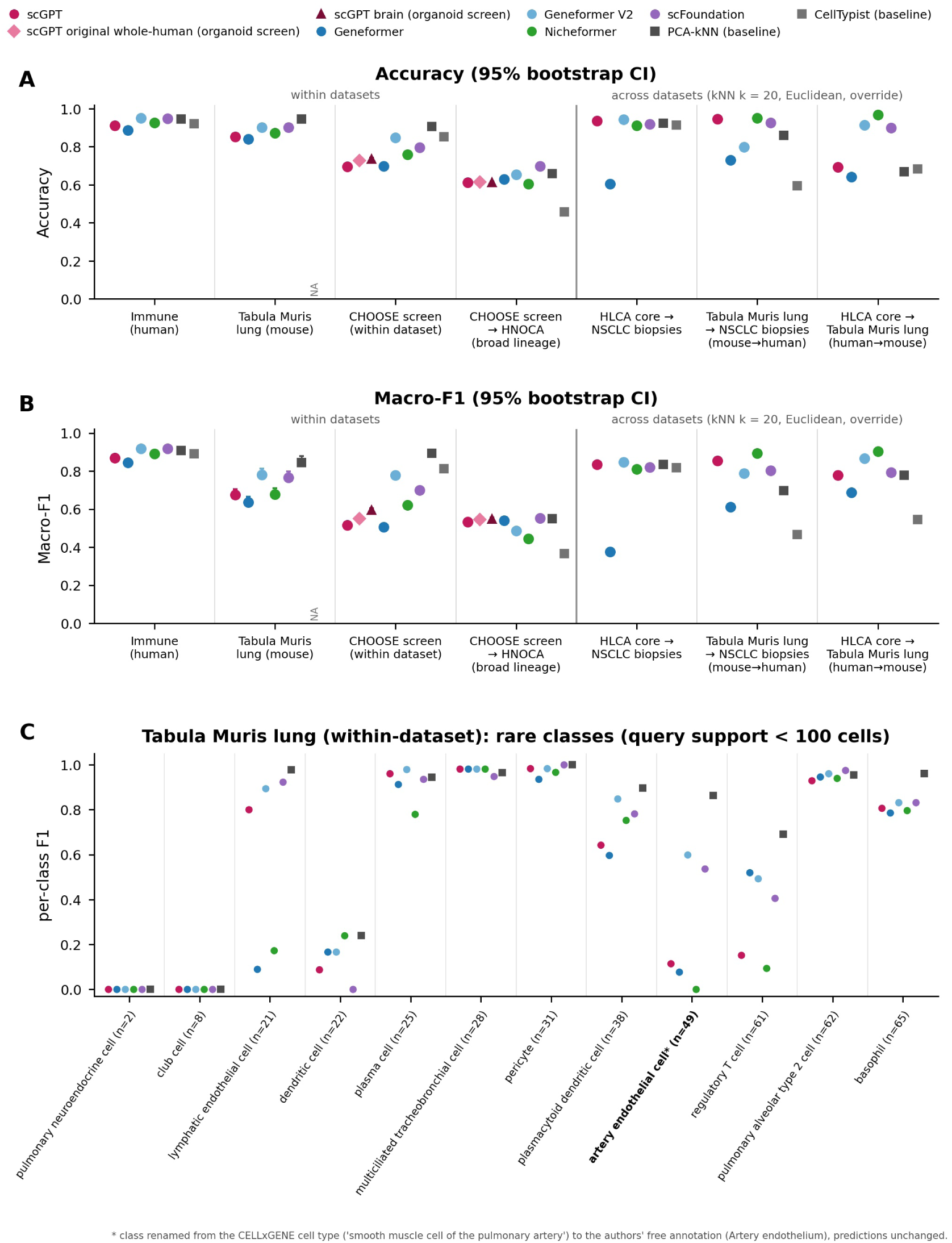
Zero-shot annotation. (A) Accuracy and (B) macro-F1 with 95% bootstrap intervals for kNN label transfer from five FM embeddings (circles) and for PCA-kNN and CellTypist baselines (squares), within datasets (Immune atlas, Tabula Muris lung, CHOOSE organoid screen cells, within the dataset and as broad-lineage concordance after HNOCA mapping) and across datasets (HLCA to lung-cancer biopsies, Tabula Muris lung to the biopsies for mouse to human, and HLCA to Tabula Muris lung for human to mouse, all with k = 20, Euclidean distance, override and coarse crosswalk). For the organoid screen the original whole-human and brain scGPT checkpoints are also shown, and the main scGPT points use the continual-pretrained whole-human checkpoint. (C) Per-class F1 for rare Tabula Muris lung classes (< 100 query cells, all classes in Supplementary Figure S1). Class names follow the authors’ annotation where the CELLxGENE label is wrong (*).

### Annotation of organoid screen cells: principal components remain the reference

On the 28,344 cells of the CHOOSE organoid screen (16 cell types), no FM matched PCA in zero-shot annotation within the dataset: PCA with kNN reached accuracy 0.906 and macro-F1 0.892, CellTypist 0.852 and 0.812, and the best FM, Geneformer V2, 0.846 and 0.778, followed by scFoundation (0.796 and 0.700) and Nicheformer (0.760 and 0.620, Figure 2, Supplementary Table S18). Among the scGPT checkpoints the brain checkpoint was the best (macro-F1 0.594, against 0.550 for the original and 0.515 for the continual-pretrained whole-human checkpoint), about 0.30 below PCA. Mapped onto HNOCA and scored as broad-lineage concordance, all embeddings reached accuracies of 0.605 to 0.698. scFoundation was the most accurate (0.698 against 0.658 for PCA) with a macro-F1 equal to that of PCA (difference +0.002, 95% CI −0.004 to 0.008), and the ranking changed when the HNOCA label Glioblast was accepted for radial glia (Geneformer V2 0.850, PCA 0.731). Fine-tuning on 4,000 cells improved every model over its own zero-shot embedding (macro-F1 gain 0.046 to 0.310), but for all five models the arm selected on validation stayed below PCA-kNN trained on the same cells (macro-F1 0.885, differences −0.041 to −0.083, all q < 0.05, Supplementary Figure S3, Supplementary Table S19).

### Perturbation in cell lines: zero-shot deletions do not predict effect size

For every representable single-gene perturbation of Norman and Adamson (75 to 152 per dataset), the predicted deletion magnitude was unrelated to the real effect size: Spearman’s ρ between predicted cosine shift and the number of real DEGs ranged from −0.08 to 0.06 for the four FMs, none significant, and ranking perturbations by target expression did no better (ρ = −0.03 and 0.06, Figure 3A). The direction of the shift was more informative. A deletion should move cells away from a CRISPRa response: Geneformer V2 and scFoundation did so for 76% and 75% of Norman perturbations (both p < 1e-8). For Adamson knockdowns scGPT (70%, p = 0.0009) and Nicheformer (71%, p = 0.0002) shifted cells in the real direction (Figure 3B, Supplementary Tables S20 to S25). With Geneformer’s own protocol, which models CRISPRa as overexpression, Geneformer V2 predicted Norman effect sizes (ρ = 0.30, p = 0.0002, BH q = 0.001) whereas V1 did not (ρ = −0.04, Figure 3A, Supplementary Table S26).

**Figure 3.**
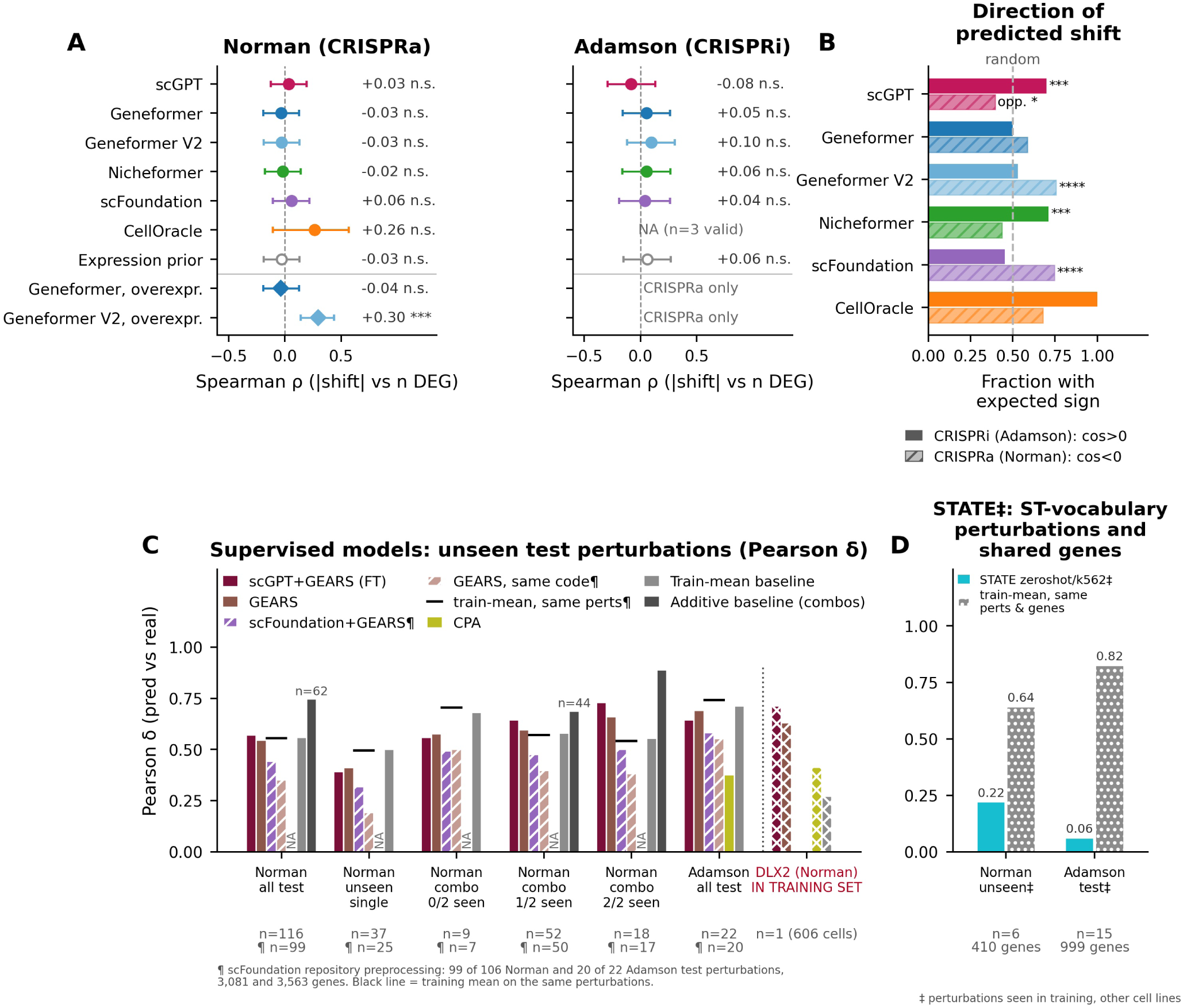
Perturbation in K562. (A) Spearman correlation between zero-shot predicted knockout magnitude and the real number of differentially expressed genes (Norman CRISPRa, Adamson CRISPRi), with the target-expression prior as baseline. The two lower Norman rows use Geneformer’s overexpression protocol, which matches the CRISPRa design. Bars, approximate 95% intervals (Fisher z). (B) Fraction of perturbations whose predicted embedding shift has the expected sign relative to the real shift (opposite for CRISPRa, where a deletion is simulated, same for CRISPRi). Asterisks, sign test, and opp., significant in the opposite direction. (C) Pearson δ of supervised models, the training-mean and additive baselines on unseen test perturbations, by Norman subgroup and for Adamson. Hatched bars, scFoundation with GEARS (frozen scFoundation embeddings as GEARS input, authors’ code and settings) and GEARS without embedding from the same code, on the scFoundation preprocessed data (¶ 99 of 106 Norman and 20 of 22 Adamson test perturbations, 3,081 and 3,563 genes), with the training mean on the same perturbations as a black line. DLX2 is marked as part of the training set. (D) STATE (‡, K562 held out) on the test perturbations in its vocabulary (Norman 6, Adamson 15), compared with the training-mean baseline on the same perturbations and on the genes shared by STATE’s output and the GEARS gene set (410 and 999 genes).

Supervised models were compared with simple baselines on unseen test perturbations (Figure 3C, Supplementary Tables S27 to S30). On Norman, GEARS reached a mean Pearson δ of 0.543 and scGPT fine-tuned 0.568, while the mean training shift reached 0.557. For unseen single genes the baseline was higher than GEARS (0.498 vs 0.409), and for combinations of two seen genes the additive baseline was higher (0.887 vs 0.658), consistent with [7]. On Adamson, the training-mean baseline (0.711) exceeded GEARS (0.688), scGPT fine-tuned (0.642) and CPA (0.375), and beat GEARS for 19 of the 22 test perturbations (Wilcoxon p = 0.0009, Supplementary Figure S4). With frozen scFoundation embeddings as GEARS input, Pearson δ exceeded GEARS run with the same code on Norman (0.440 versus 0.349) but not the training mean (0.556), and on Adamson reached 0.580 against 0.742, on the 99 and 20 test perturbations available in its inputs (Supplementary Table S31).

STATE could be scored only on perturbations in its vocabulary, all seen as CRISPRi knockdowns in other cell lines during training. On the test perturbations it covers (6 Norman, 15 Adamson), with K562 held out, it reached a mean Pearson δ of 0.22 and 0.06, against 0.64 and 0.82 for the training mean on the same perturbations and genes (Figure 3D, Supplementary Table S32). It predicted the direction of Adamson knockdowns but not their magnitude (ρ = 0.10).

### Perturbation in neural tissue: predicted effects track target expression

In the 991 control cells of the organoid screen, the predicted effect of deleting each of the 36 genes followed the target’s expression in each cell type for every zero-shot model (ρ = 0.56 to 0.93 over 504 gene by cell-type pairs, highest for Nicheformer and Geneformer V1, Figure 4A, Supplementary Tables S33 to S35), and the same held in the mouse cortex (ρ = 0.70 to 0.97 over gene by class pairs, Supplementary Table S36). Fine-tuning as a cell-type classifier rescaled these effects (median ratio to zero-shot 0.85 to 6.24) while preserving their ranking (ρ = 0.77 to 0.99 over pairs, Figure 4B, Supplementary Table S37).

**Figure 4.**
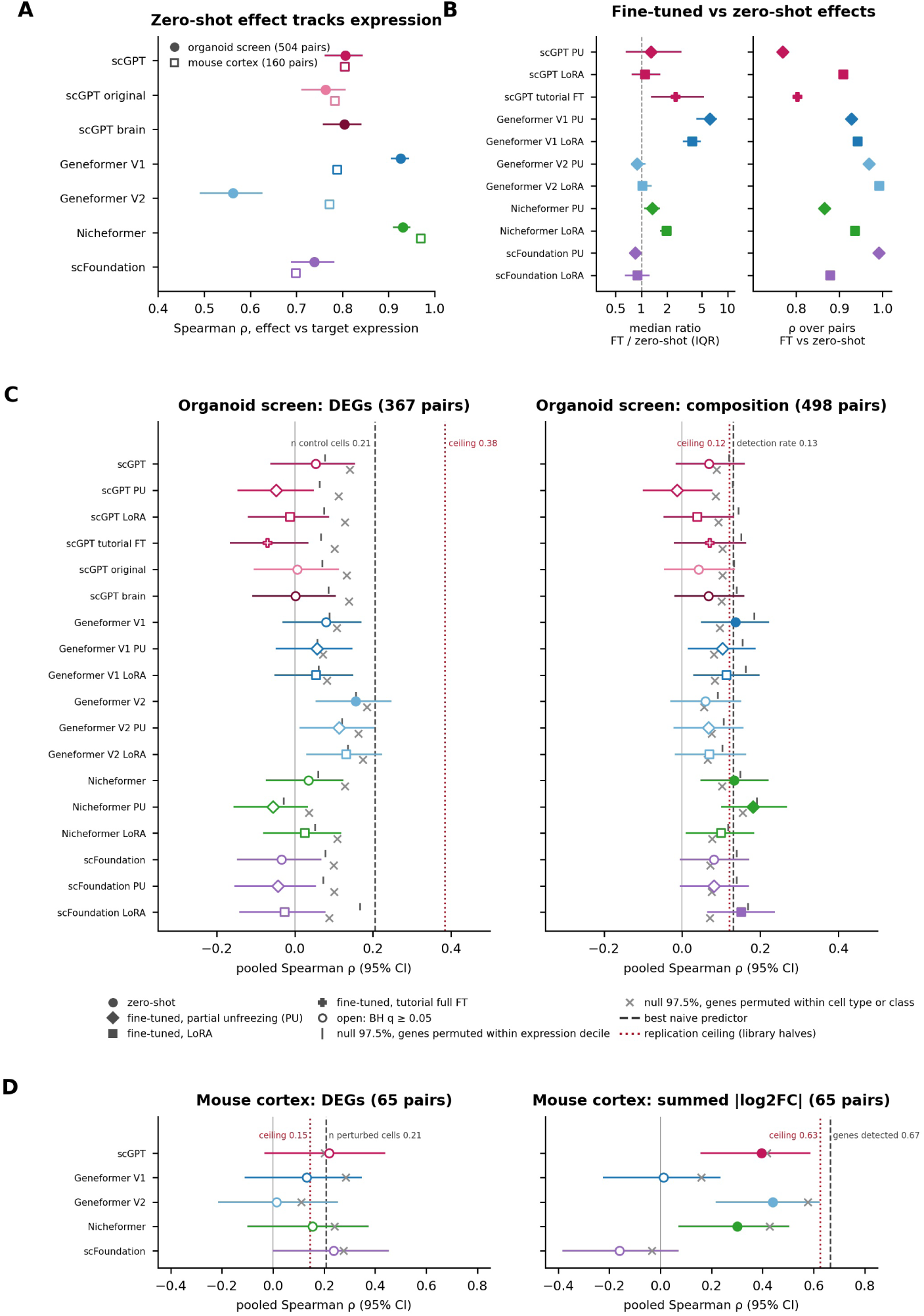
Perturbation in neural tissue. (A) Spearman ρ between each zero-shot model’s predicted per-cell-type knockout effect and target-gene mean expression in the 991 control cells of the organoid screen (36 genes, 504 gene by cell-type pairs, filled circles, 95% bootstrap CI) and in the 374 GFP control cells of the mouse cortex (160 gene by class pairs with a prediction, open squares). (B) Fine-tuned versus zero-shot knockout effects in the organoid screen for the eleven fine-tuned backbones (PU, partial unfreezing, LoRA, tutorial full fine-tuning): median ratio with interquartile range (log scale) and rank correlation over the 504 pairs. (C) Organoid screen: pooled Spearman ρ (95% CI) of zero-shot and fine-tuned predictions with the number of DEGs (367 pairs) and with the absolute composition log odds ratio (498 pairs). Filled markers, BH q < 0.05 in the family of 36 tests. Grey ticks and crosses, 97.5% quantile of the nulls that permute genes within expression decile and within cell type. Dashed line, best naive predictor. Dotted red line, split-half replication ceiling. (D) Mouse cortex: the same for the number of DEGs and the summed absolute log2 fold change against GFP control cells (65 gene by class pairs, primary family, null within cell class, ceiling between odd and even libraries).

### Measured effects in neural tissue: no model exceeds naive predictors

In the organoid screen none of the 36 tests of the predefined rule was met (18 zero-shot and fine-tuned predictors by two ground truths, Figure 4C, Supplementary Table S38). Five predictions were positive at q < 0.05, Geneformer V2 against the number of DEGs (ρ = 0.156) and Geneformer V1, Nicheformer, Nicheformer with partial unfreezing and scFoundation with LoRA against composition changes (ρ = 0.133 to 0.181), but none exceeded the best naive predictor by 0.10: the number of control cells per cell type reached ρ = 0.206 on the DEG truth and the detection rate of the target 0.130 on the composition truth. The DEG associations did not exceed a null that permutes genes within cell type, so they reflect differences between cell types, and the composition associations did not exceed a null that permutes genes within expression decile, so they reflect target expression, as does the detection rate. Under the within-cell-type null, the nominal associations of scGPT and Geneformer V2 with DEG numbers found with a per-gene Fisher combination (p = 0.048 and 0.020, Supplementary Table S39) were not above chance (permutation p = 0.26 and 0.45, Supplementary Figure S7C, Supplementary Table S40). Fine-tuning did not improve agreement with the DEG truth (change −0.123 to +0.009, interval below 0 for 6 of 11 backbones and never above 0). The measured effects themselves replicated only partly between halves of the libraries (ρ = 0.38 for DEG numbers and 0.12 for composition, Supplementary Table S4).

In the mouse cortex none of the 10 primary tests (five models by two measures of effect, 65 gene by class pairs) met the rule (Figure 4D, Supplementary Table S41). Against the number of DEGs the highest correlations, scFoundation (ρ = 0.24, q = 0.14) and scGPT (0.22, q = 0.16), did not exceed the number of perturbed cells alone (0.21). Against the summed fold change, Geneformer V2 (ρ = 0.44, q = 0.002) and scGPT (0.40, q = 0.005) were below the genes detected per cell class (0.67) and were reproduced by permuting genes within cell class (null mean 0.47 and 0.26). With 374 control cells the number of DEGs agreed poorly between odd and even libraries (ρ = 0.15, Supplementary Figure S7A, Supplementary Table S42), so this screen sets a low ceiling for any predictor, although its positive controls were reproduced, including the composition effect of Dyrk1a (fewer microglia, log odds ratio −0.92, FDR 0.002). Nicheformer with the mouse token behaved like the human-trained models with orthologs (ρ = 0.15 and 0.30), and the secondary reference gave the same result (Supplementary Figure S7B, Supplementary Table S43). On the 14 genes shared with the organoid screen the measured burden agreed only weakly between the two systems (ρ = 0.60, 95% interval −0.04 to 0.94, secondary reference), and no model was consistent across them (Supplementary Table S44).

### A larger model version improves annotation but not perturbation

We repeated every Geneformer row with the V2-316M release (Supplementary Table S22). V2 annotated better than V1 on immune cells, Tabula Muris lung, the biopsies, the organoid screen (macro-F1 0.778 versus 0.504) and from human to mouse (0.914 versus 0.642). Zero-shot deletion results did not change: V2 tracked neither the organoid screen nor the mouse cortex effects beyond naive predictors, and its network stayed near chance against K562 knockdown targets (FE@50 1.02). Scale therefore helped annotation and, with the overexpression protocol, CRISPRa effect sizes.

### Knockout effects are mostly non-specific

The expression-matched specificity test explains why predicted effects follow target expression (Figure 5, Supplementary Table S6). In the organoid screen, deleting the target shifted the embedding no more than deleting one of 20 control genes matched for expression in the same cell: the target exceeded all 20 controls in 1.1% to 5.7% of gene by cell-type pairs, against 4.8% expected by chance (binomial p ≥ 0.19), and the fraction of pairs with z > 2 was 0.9% to 6.6%. The median ratio of target to matched-control effect was close to 1 for the rank-encoded models (Geneformer V1 and V2 0.98, Nicheformer 0.99) and below 1 for the value-encoded models (scGPT 0.49 to 0.56, scFoundation 0.28). Once the matched null was subtracted, the correlation of the effect with target expression disappeared or became negative (ρ −0.73 to −0.04).

**Figure 5.**
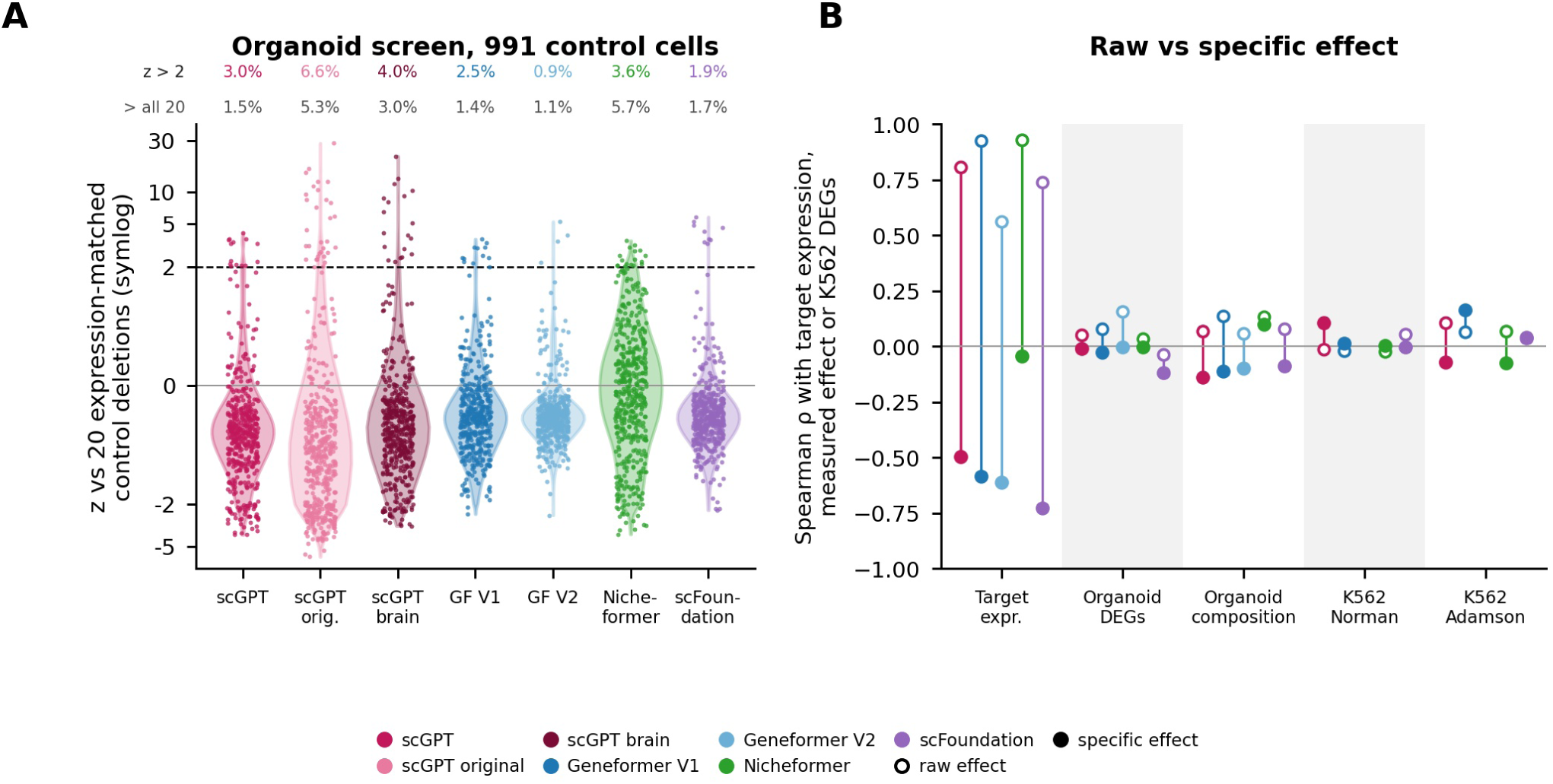
Expression-matched specificity test for in silico knockout. (A) Distribution of z (observed shift minus the mean of 20 expression-matched control deletions, divided by their standard deviation) per gene by cell-type pair in the 991 control cells of the organoid screen, for the seven zero-shot models and checkpoints, with the fraction of pairs above z = 2 and the fraction in which the target exceeds all 20 controls (4.8% expected by chance). Controls were computed on all expressing cells, and on at most 200 expressing cells per gene for Geneformer V2 and scFoundation. (B) Spearman ρ of raw (open) and specific (filled) effects with target-gene expression, with the measured effects of the organoid screen (number of DEGs, composition) and with the number of differentially expressed genes per perturbation in K562 (Norman, Adamson, not run for Geneformer V2).

Removing the non-specific component did not improve agreement with real knockouts. In K562, the specific effect correlated with the number of DEGs or the energy distance at ρ −0.16 to 0.17 (all p ≥ 0.13, Supplementary Table S45). In the organoid screen only the specific effect of Nicheformer correlated positively with composition changes (ρ = 0.10, p = 0.027), below the detection rate of the target, and no specific effect correlated with DEG numbers.

### Gene regulatory networks: co-expression is a strong baseline

Across K562 and biopsies, FM networks recovered known regulatory targets close to chance (Figure 6, Supplementary Tables S46 to S49). For Adamson knockdown targets, the best expression-matched FE@50 was 1.32 (scFoundation attention), equal to the expression prior (1.31) and below co-expression (1.61). For K562 ChIP-seq targets the best FM network (scGPT attention, 2.20) was also below co-expression (2.87). In the biopsies, the best FM network (scGPT embedding, 1.41) matched co-expression (1.40), and with patients as the unit of analysis (pseudobulk, nominal p < 0.01) the expression-matched Geneformer V2 network exceeded it (FE@50 2.90 versus 2.04), although on few hubs (Supplementary Figure S5, Supplementary Table S50). Fine-tuning did not change this: networks from backbones fine-tuned for perturbation or K562 perturbation identity did not exceed co-expression on any primary ground truth, and perturbation fine-tuning lowered scGPT attention enrichment for knockdown targets (0.62 on the two held-out Adamson hubs, zero-shot 2.04, Supplementary Table S51).

**Figure 6.**
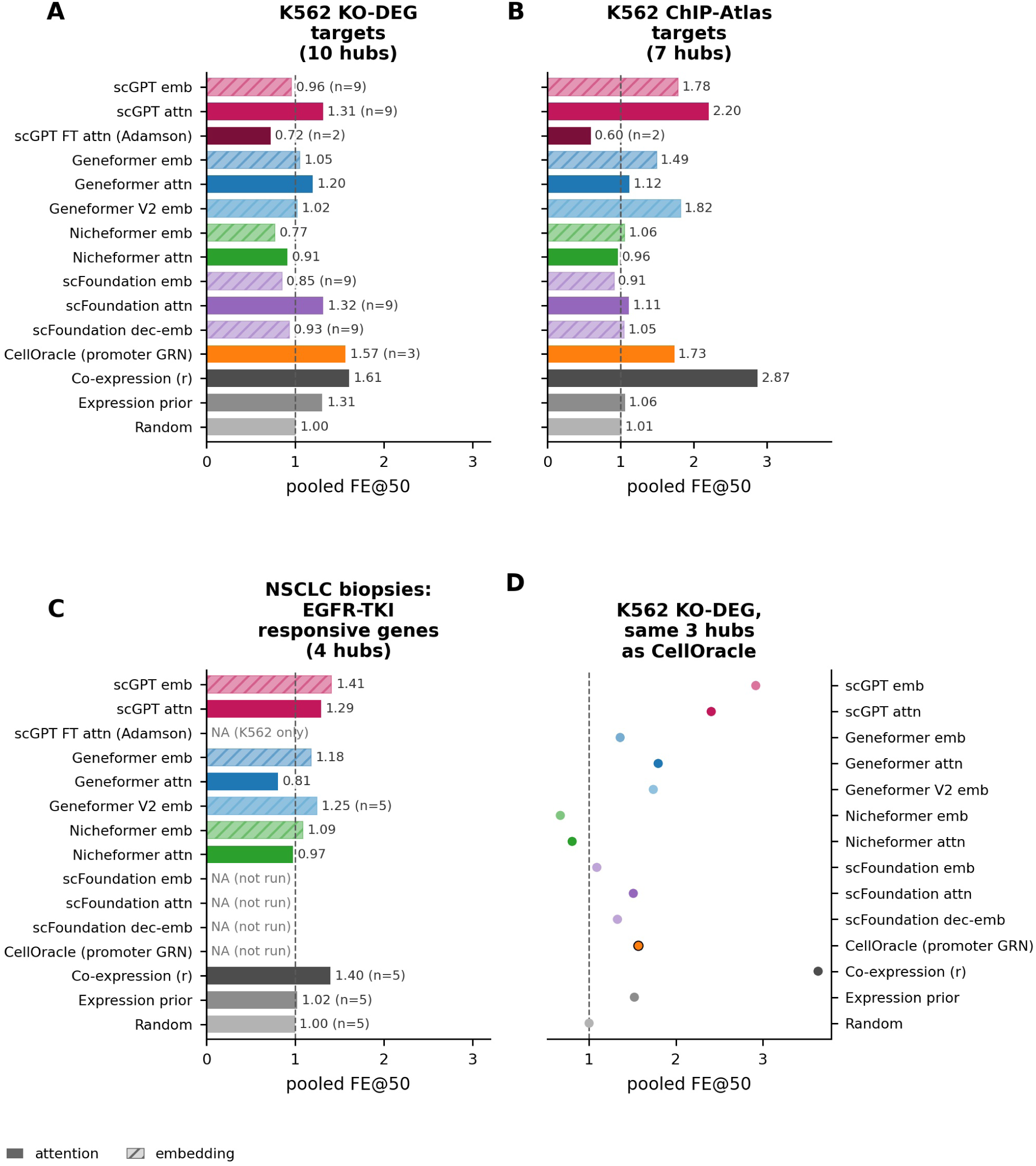
Gene regulatory networks. Pooled expression-matched fold enrichment of the top-50 predicted targets per hub (FE@50, dashed line at chance) for FM embedding (hatched) and attention (solid) networks, CellOracle, co-expression, an expression prior and random ranking, against (A) K562 knockdown targets, (B) K562 ChIP-seq targets, (C) genes responding to EGFR inhibition in treatment-naive cancer cells of the lung-cancer biopsies. (D) K562 knockdown targets for the three hubs CellOracle can score. NA, not computed.

### Supervised models trained on the organoid screen

Trained on the organoid screen itself with five-fold gene hold-out, GEARS, scFoundation with GEARS and scGPT fine-tuned predicted held-out genes (Pearson δ 0.27, 0.33 and 0.54 on the top 20 DE genes) below the mean training response (0.68, Supplementary Table S52), and Geneformer fine-tuned as a perturbation classifier did not beat its zero-shot version (median ρ −0.05 versus 0.05 for V1, 0.20 versus 0.28 for V2) or a permuted-label control (0.20 for V2, Supplementary Table S53). CellOracle, which can simulate six of the 36 genes, correlated with composition changes over 84 gene by cell-type pairs (ρ = 0.43, permutation p = 0.001) but not more than the detection rate of the target on the same pairs (0.36, Supplementary Table S40). The earlier per-gene analysis of the screen is given in Supplementary Figure S6 and Supplementary Tables S54 to S64.

### Human EGFR-TKI lung cancer: patient effects dominate cell-level ground truth

In biopsies of EGFR-mutant lung cancer before targeted therapy (TN, 7 patients) and at residual disease (RD, 8 patients) [28], every method placed the cancer compartment first after an in silico EGFR knockout, and so did the naive EGFR-expression predictors (Figure 7, Supplementary Tables S65 to S68). Against cell-level RD-versus-TN DEG counts all methods correlated positively (ρ = 0.20 to 0.53), but the progression contrast (PD versus TN) gave equal values for scGPT (0.52) and CellOracle (0.53). With patients as the unit (pseudobulk), seven of eight methods correlated negatively with the ground truth. The cell-level agreement thus reflects differences between patients and biopsies rather than treatment (Supplementary Figure S5).

**Figure 7.**
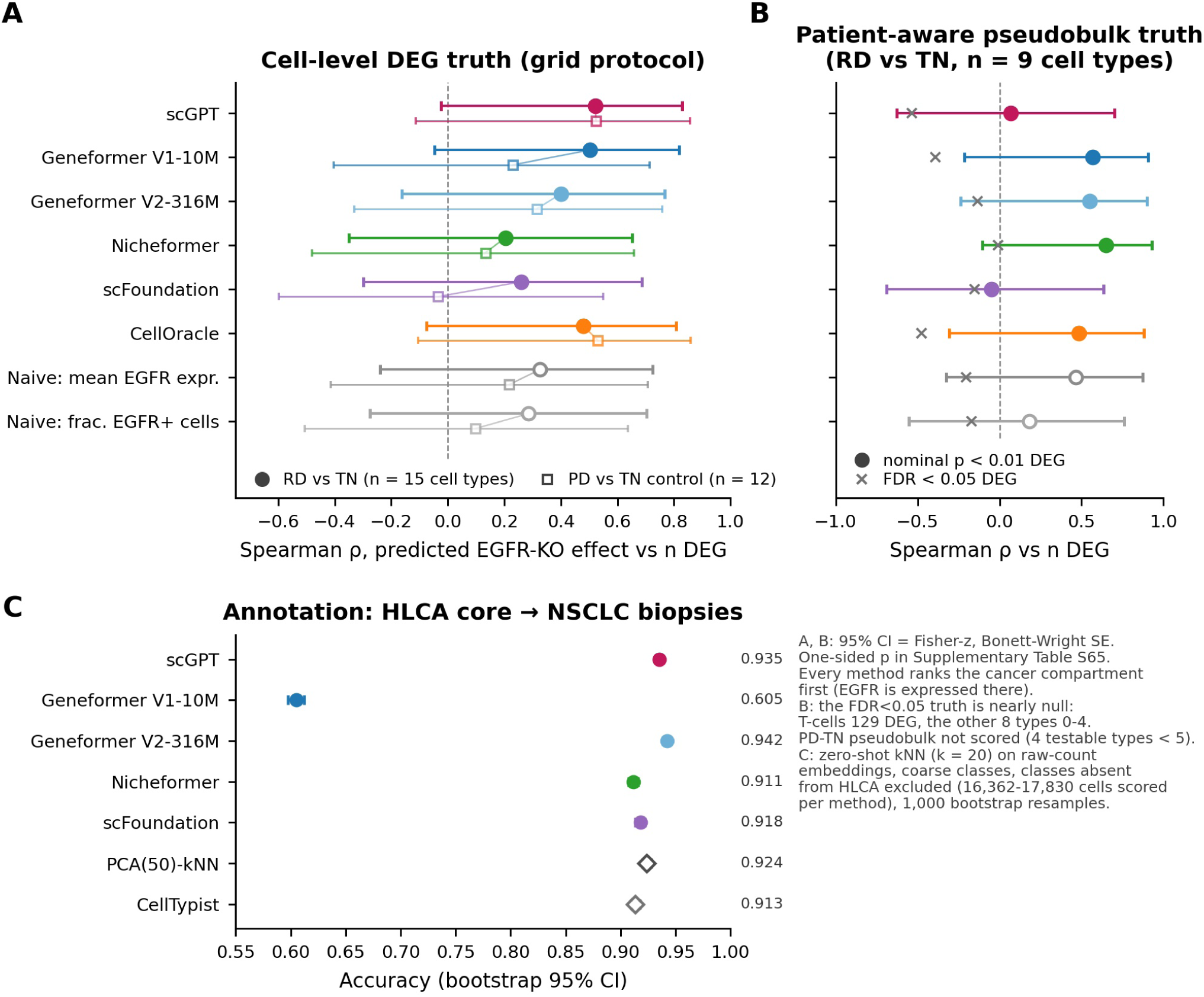
EGFR-mutant lung-cancer biopsies. (A) Spearman ρ between the predicted EGFR-knockout effect per cell type and the number of genes differentially expressed at residual disease versus before therapy, with the progression contrast as control (paired markers). (B) The same against the patient-aware pseudobulk ground truth. (C) Annotation accuracy from the HLCA reference for FMs and baselines. Intervals are approximate 95% CIs.

### Genome-scale screens: measured effects replicate but are not predicted

In the genome-scale CRISPRi screen of primary CD4 T cells from four donors [14], zero-shot effects of scGPT, Geneformer V1 and V2 and Nicheformer correlated weakly with the authors’ number of DEGs at rest (ρ = 0.091 to 0.114, BH q 0.016 to 0.050, 593 to 600 targets, Figure 8A), and scFoundation did not (ρ = 0.040). None exceeded the detection rate of the target in the same control cells (ρ = 0.14), none survived the permutation within expression decile (permutation p 0.53 to 0.99), and no model met the rule in any of the 15 tests of five models by three culture conditions (Figure 8B, Supplementary Tables S69 and S70). The measured effects themselves agreed only moderately between donors (mean ρ 0.22 to 0.26), and the predictions agreed with the per-donor effects at mean ρ 0.04 to 0.10 (Supplementary Figure S8, Supplementary Table S71). The predicted shift was no closer to the real shift of the knocked-down cells than to the shifts of other targets (median difference −0.008 to 0.009), and deleting the target moved the embedding more than 20 expression-matched control genes (z > 2) for at most 6.2% of targets, against 2.3% expected (Supplementary Tables S72 and S73).

**Figure 8.**
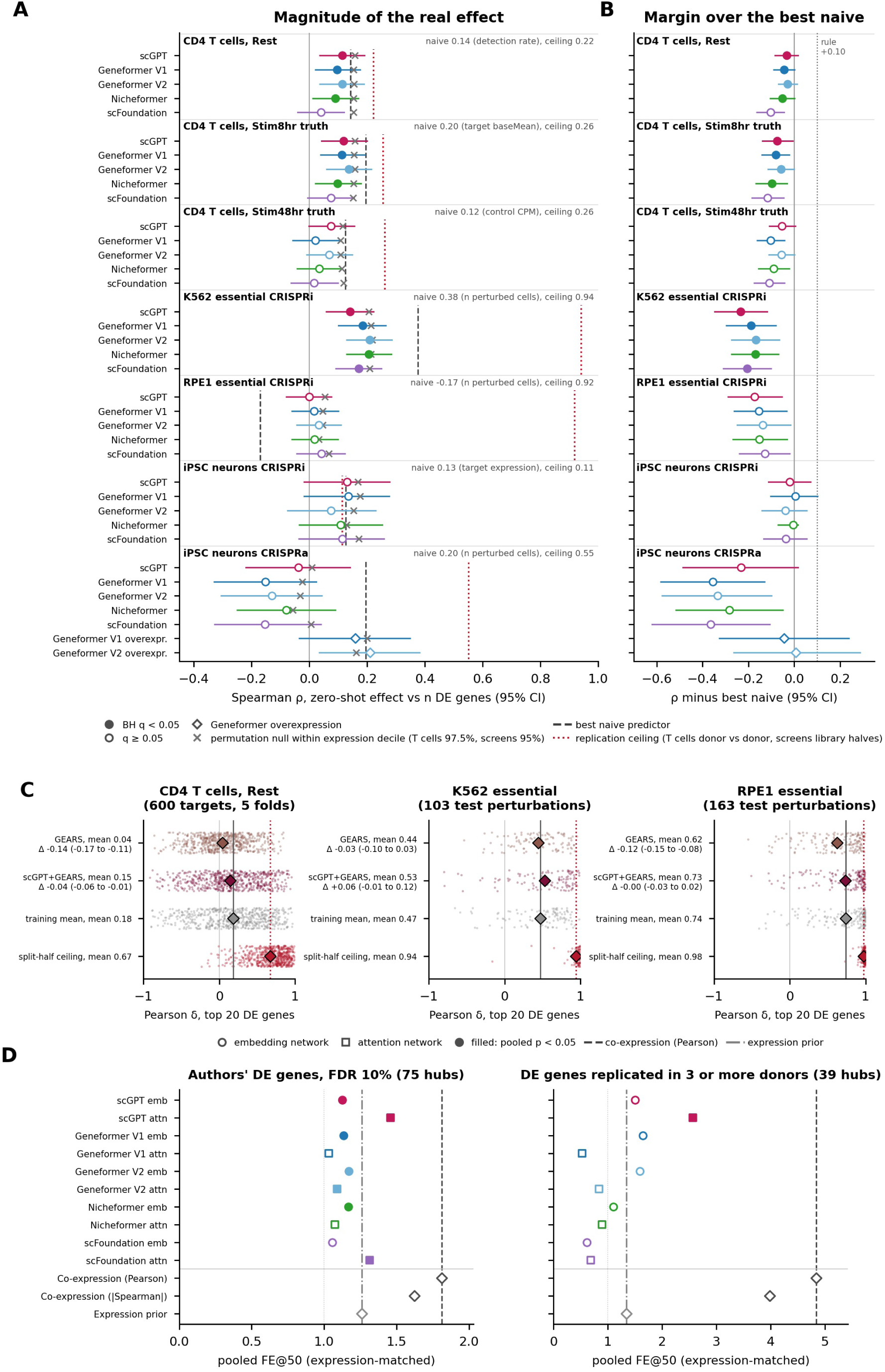
Genome-scale perturbation screens. (A) Spearman ρ (95% bootstrap CI) between the zero-shot deletion effect and the real number of DE genes, per model and screen. For primary CD4 T cells, the predictions come from 200 resting non-targeting control cells and are scored against the authors’ DE genes in resting cells and, with the same predictions, against the 8 h and 48 h stimulation truths (593 to 600 targets). The other screens are K562 and RPE1 essential-gene CRISPRi (the same 600 targets in both), iPSC-derived neuron CRISPRi (176 to 180 targets) and CRISPRa (96 or 97 targets, plus Geneformer overexpression, diamonds). Filled markers are BH q < 0.05 within the pre-registered families (15 T cell tests, 20 scPerturb tests, 4 overexpression tests). Crosses are the permutation null with real effects permuted within expression deciles (97.5% quantile for T cells, 95% quantile for the scPerturb screens, as computed by each track). Dashed lines are the best naive predictor on all targets (value and predictor in the group header). Red dotted lines are the replication ceiling. For T cells this is the mean donor-versus-donor ρ of the per-donor edgeR recomputation (6 donor pairs). For the scPerturb screens it is library half A against half B. (B) ρ minus the best naive predictor on the same targets, with the paired bootstrap 95% CI. For the scPerturb screens the absolute ρ of the best naive predictor is subtracted. The dotted line marks the margin of 0.10 required by the decision rule. No model meets the rule in any test (0 of 15 T cell tests, 0 of 20 scPerturb tests). (C) Supervised models. Points are the Pearson δ on the top 20 DE genes of each held-out target (T cells, five-fold target hold-out over 600 targets in resting cells) or test perturbation (K562 and RPE1, GEARS simulation split, 103 and 163 test perturbations). They are shown for GEARS, scGPT with GEARS, the training-mean prediction and the split-half ceiling. Diamonds are means. The labels give the mean and the mean difference from the training mean with its 95% CI. The solid grey line is the training mean and the red dotted line the split-half ceiling. (D) T cell networks. Pooled expression-matched fold enrichment of the top 50 predicted targets (FE@50) of transcription-factor hubs, for embedding (circles) and attention (squares) networks extracted from 2,000 resting non-targeting cells. Filled markers are pooled p < 0.05. The dashed line is signed Pearson co-expression on the same cells and the dash-dot line the expression prior. Absolute Spearman co-expression is shown as a sensitivity reference. Left: authors’ DE genes at 10% FDR (75 hubs, 71 to 75 per model). Right: DE genes replicated in at least 3 donors (39 hubs).

In four screens of the scPerturb collection [25], the essential-gene CRISPRi screens of K562 and RPE1 [13] with the same 600 targets and the CRISPRi and CRISPRa screens of iPSC-derived neurons [17] with all 184 and 100 targets, no model met the rule in any of 20 tests (Figure 8A,B, Supplementary Tables S74 and S75). In K562 all five models correlated with the number of DEGs (ρ = 0.14 to 0.21, q ≤ 0.002), as did the expression of the target (0.21), and the number of perturbed cells reached 0.38. In RPE1 the correlations were close to zero (ρ = 0.00 to 0.04). These measured effects replicated almost perfectly between halves of the libraries (ρ = 0.94 in K562 and 0.92 in RPE1, Supplementary Table S76), so the failure does not come from a noisy ground truth, unlike in the neuron CRISPRi screen, whose effects did not replicate (ρ = 0.12). The part of the effect specific to the cell line, K562 minus RPE1, was also reproducible (ρ = 0.85) but was predicted by no model (ρ = −0.085 to 0.024, Supplementary Figure S8, Supplementary Table S75). Predicted shifts agreed in sign with the real shifts for most targets in 11 of 20 model by screen pairs, but pairing each prediction with the real shift of another target gave nearly the same agreement (excess at most 0.063, Supplementary Table S77). On the CRISPRa neuron screen Geneformer’s overexpression protocol gave ρ = 0.16 to 0.23 (q ≥ 0.076), not above the number of perturbed cells (0.20, Supplementary Table S78).

Supervised models did not beat the training mean on these screens (Figure 8C, Supplementary Tables S79 and S80). In T cells, with five-fold target hold-out, GEARS and scGPT with GEARS reached a Pearson δ of 0.041 and 0.146 against 0.182 for the training mean (split-half ceiling 0.672). With the GEARS split, GEARS was below the training mean in K562 (0.442 against 0.473) and RPE1 (0.622 against 0.738), and scGPT with GEARS was at its level (0.530 against 0.473 in K562, 95% interval of the difference −0.011 to 0.125, and 0.734 against 0.738 in RPE1). FM networks recovered the knockdown targets of up to 75 T cell transcription factors less well than co-expression on the same cells (best FM, scGPT attention, FE@50 1.46 against 1.81, Figure 8D, Supplementary Table S81).

### Gene representations of FMs predict which knockdowns have large effects

Because deleting a gene from the input did not reveal how strongly its knockdown changes the cell, we asked whether the gene representations that FMs learn during pretraining carry this information. Two re-readings of the stored deletion effects did not help. Dividing the effect by the fraction of control cells that express the target, which removes its dependence on detection, lowered the correlation in T cells (ρ = −0.03 to 0.06) and met the rule in 0 of 15 T cell and 0 of 20 cell-line tests, and adding the deletion effect to expression, detection and number of perturbed cells improved the out-of-fold prediction in none of 35 tests (Supplementary Tables S82 and S83). Simulating a partial knockdown matched to the residual expression measured in each screen, instead of a deletion, did not change this: it met the rule in 0 of 15 T cell and 0 of 15 cell-line tests. Added to the naive predictors, it gave small gains in 2 of 15 cell-line tests (Geneformer V2 in K562, 0.048, q = 0.037, and scGPT in neuron CRISPRi, 0.054, q = 0.0498), which disappeared with a fixed residual expression of 0.2. Compared with deletion on the same targets, knockdown did not change ρ in T cells, lowered it for scGPT and scFoundation in K562 (by 0.12 for both) and raised it by 0.01 for Geneformer V1 in K562 and RPE1 (Supplementary Methods). Gene vectors trained on one system did not transfer to another: from K562 to RPE1, from RPE1 to K562 and from either to T cells no FM exceeded expression or STRING degree (0 of 20 tests, for example ρ = 0.20 for scGPT against 0.37 for STRING degree from RPE1 to K562, Supplementary Table S84).

Within a system, however, adding the gene vectors of scGPT or Nicheformer to expression features improved the prediction of the number of DEGs in K562 (ρ from 0.39 to 0.46 and from 0.40 to 0.46) and in T cells (from 0.20 to 0.25 and to 0.24), above 200 residual-permutation nulls (p = 0.005, the smallest attainable, Supplementary Figure S9, Supplementary Tables S85 and S86). We tested this observation on an independent screen with a plan written before its results. In the genome-wide CRISPRi screen of K562 [13], with 7,509 and 7,557 targets never used in this study, the gene vectors of scGPT and Nicheformer raised the correlation from 0.26 to 0.43 and 0.41 (gain 0.17 and 0.15, 95% intervals 0.15 to 0.19 and 0.13 to 0.17, p = 0.005, q = 0.005), whereas the 95th percentile of the gains on 100 fully random truths was 0.05. Against a nonlinear baseline that also included STRING degree, an analysis added after the primary results, the gains were 0.10 and 0.09 (both p = 0.005, Supplementary Figure S9, Supplementary Tables S86 and S87). The same protocol, applied unchanged to the other three models, confirmed the gain for Geneformer V2 and V1 (from 0.26 to 0.37 and 0.34, gain 0.110 and 0.086, q = 0.0075 for both, and 0.063 and 0.051 against the nonlinear baseline) but not for scFoundation (gain −0.007, q = 0.12), so the five models ranked scGPT, Nicheformer, Geneformer V2, Geneformer V1 and scFoundation (Supplementary Table S88).

We also asked, with a plan written before the results, whether in the genome-scale screens the zero-shot deletion shift points towards the real response of the deleted gene more than towards the responses of other targets, and more than the shifts predicted for 20 expression-matched control genes deleted in the same cells. Of 20 tests (five models, four screens), three were positive against other targets and two also against the matched controls (Nicheformer in K562 and Geneformer V2 in RPE1), with a median gain in cosine of about 0.013, roughly 4% of the agreement between two halves of the same perturbed cells (Supplementary Table S89). No model passed in the T cells or in the iPSC neurons, and only the Geneformer V2 result in RPE1 held in every sensitivity analysis, so in these screens the deletion shifts carried at most a faint gene-specific direction.

Four newer public models, ranked before any score by the number of benchmarks that include them, behaved like the other FMs (Supplementary Table S90). From human to mouse UCE, SCimilarity, CellPLM and TranscriptFormer exceeded CellTypist with default settings (0.81 to 0.98 against 0.68), but only UCE (0.979) also exceeded tuned CellTypist (0.941). From mouse to human UCE and SCimilarity exceeded PCA by 0.09 and 0.10, a margin that fell below 0.01 with cosine kNN. Within a dataset none matched PCA. Their gene vectors, however, added to expression features on the genome-wide K562 screen (ρ between 0.39 and 0.44, against 0.26 with expression alone, all above the Freedman-Lane null), whereas their deletions did not beat the number of perturbed cells.

## Discussion

Evaluated against simple baselines on the same data, current single-cell FMs helped in specific settings: mapping a reference across species (also for newer models such as UCE), the direction of perturbation responses in K562 (only faintly in genome-scale screens), CRISPRa effect sizes with Geneformer’s own overexpression protocol and, with patient-level ground truth, tumour regulatory networks. Within a species their embeddings support label transfer about as well as principal components, as reported for zero-shot clustering [5,6], although fine-tuned Geneformer V2 exceeded them on Tabula Muris lung, and on organoid screen cells principal components stayed ahead also after fine-tuning. Across species the margin over CellTypist depended on tuning its regularisation, which a fair benchmark must do. Their zero-shot in silico knockouts do not predict how strongly a real knockout perturbs a cell line and, in neural tissue, largely restate where the target gene is expressed. Their gene-gene networks do not recover knockdown targets or ChIP-seq binding better than co-expression. These results extend earlier conclusions from cell lines [5,6,7,8] to genome-scale screens in primary T cells and two cell lines, an organoid screen, an in vivo screen and tumour biopsies.

Perturbation screens in tissue provide a measured effect for every gene and cell type, but in both tissue screens analysed here the measured effect replicated only partly between libraries, and naive predictors that know nothing about gene function, the number of perturbed or control cells and the detection rate of the target, reached the correlations of the best models. In the K562 and RPE1 screens, by contrast, the measured effects and their difference between the two cell lines replicated almost perfectly between libraries, and no model predicted them either, so a noisy ground truth does not explain the failure. Associations found with per-gene statistics disappeared under a null that permutes genes within cell type, which should therefore accompany any such score. Agreement between methods was here mostly shared dependence on target expression, which a naive predictor reproduces.

The specificity test gives a mechanism for this dependence and a practical control. Deleting a gene from a cell’s input changes the embedding roughly as much as deleting any other gene of similar expression, so the per-cell-type effect mostly measures how many cells express the target. In the organoid screen the target exceeded its matched controls no more often than expected by chance, and the excess did not predict measured effects. We propose that in silico perturbation results be reported against such a null. The test is implemented in fmaudit.

Fine-tuning did not change this picture. It improved classification over each model’s own embedding without reaching principal components within a species, rescaled predicted effects up to six-fold without changing their ranking, and did not yield better gene networks, so it altered sensitivity rather than correctness. Supervised models trained on Perturb-seq of cell lines and primary T cells or on the organoid screen itself, including GEARS with scFoundation embeddings, did not beat the mean training shift for unseen genes and cannot be applied to systems without perturbation data.

Without the checks applied here, a fine-tuning run that silently fell back to the pretrained model would have suggested that fine-tuning leaves predictions unchanged. An input re-sampled before and after each deletion would have made scGPT’s effects unrelated to the deleted gene. A cell-level ground truth would have rewarded predictors of cell number, and a per-gene statistic without a within-cell-type null would have rewarded differences between cell types. Running Nicheformer without its species and assay tokens changed its values, although not the conclusions (Supplementary Table S91). Table 1 summarises the resulting recommendations, including a target-detection check now in fmaudit.

**Table 1.** Practical guidance for applying single-cell foundation models to new data.

| When | Recommendation |
|---|---|
| Annotating cells within one species | Start with label transfer on principal components or with CellTypist. In this benchmark zero-shot FM embeddings did not do better. |
| Transferring labels between species | A model pre-trained on both species may help. Compare it with a principal-component or highly-variable-gene baseline on the same cells, with both Euclidean and cosine classifiers. |
| Before simulating a knockout | Check that the target is detected in enough cells of the cell types of interest. A deletion cannot act where the gene is absent from the input, and it represents neither effects on other cells nor missense variants. |
| Interpreting knockout predictions | Compare them with a ranking of cell types by target expression and with deletions of expression-matched control genes. Agreement between models is not validation. |
| Validating against real perturbations | Use libraries or donors, not cells, as replicates, report the agreement between replicates as a ceiling, and compare scores with a null that permutes genes within cell type. Per-cell DEG counts follow the number of cells. |
| Predicting the size of a perturbation response | Zero-shot deletions did not predict it, and a model-native overexpression protocol did so only modestly. The gene vectors of most FMs (seven of eight tested, not scFoundation), used as features of a simple regressor trained on measured effects of the same system, added to expression features (and, in a robustness analysis, to STRING degree) but did not transfer between systems. Supervised models need perturbation data from the same system and should be compared with the mean training response. |
| Inferring regulatory networks | Compare FM-derived networks with co-expression, which they did not exceed on any primary ground truth. |
| Before trusting any result | Check the input scale of every matrix, confirm that fine-tuning changed the weights and keep test perturbations out of training. |

Limitations: the mouse cortex screen has only 374 control cells and measured effects that replicate poorly between libraries, and the organoid screen has a modest replication ceiling, so both test the models against a noisy truth. Three mouse targets had no human ortholog in our table. In silico deletion represents only cell-autonomous effects, and FMs see only expression, so missense variants will require models that combine expression with sequence. In the T cell screen only resting cells were embedded, so the stimulated conditions test resting predictions, and the two slowest models were run on 100 control cells in three screens. The gene-vector gains were measured within a system and did not transfer between cell lines or to T cells. The biopsies come from few patients. Newer models may behave differently.

## Conclusions

Across three tasks, from cell lines and primary T cells to an organoid screen, an in vivo screen and tumour biopsies, single-cell FMs were matched by simple baselines in most comparisons, and helped mainly to transfer labels across species and, in K562, to give the direction of perturbation responses. No model predicted measured knockout effects beyond naive predictors and permutation nulls, in neural tissue or in genome-scale screens whose effects replicate. The gene vectors learned in pretraining by seven of the eight models tested, in contrast, predicted which knockdowns have large effects beyond expression, confirmed on an independent screen, which suggests using FMs as a source of gene features rather than as simulators of deletions. Task-matched baselines, these nulls, replication ceilings and pipeline checks should become a standard component of FM benchmarks.

## Ethics Statement

This study used only public datasets, according to their original consent and data-use terms.

## Data Availability

Public data were obtained from their original sources: Norman K562 Perturb-seq (GEO GSE133344) and Adamson K562 Perturb-seq (GEO GSE90546), both in GEARS processing, the K562 and RPE1 essential-gene screens and the iPSC-derived neuron screens from the scPerturb collection (Zenodo record 13350497) [25], with the GEARS-processed essential-gene files from Harvard Dataverse, the primary CD4 T cell Perturb-seq from the public release of its authors on the CZI virtual cell models platform [14], the Immune Human atlas from the scIB benchmark [23], HNOCA [26], Tabula Muris Senis (GEO GSE132042) [24], the CHOOSE screen from Zenodo record 7083558 [15], the in vivo Perturb-seq of the mouse cortex from GEO GSE157977 [16] with cell annotations from Single Cell Portal study SCP1184 [27], the EGFR-mutant lung-cancer biopsies from NCBI BioProject PRJNA591860 [28] and the HLCA core from CELLxGENE [29]. Code, configurations, SLURM scripts, environment specifications, the analysis plans and the result tables underlying every figure and Supplementary Table (without per-cell arrays, embeddings and model checkpoints) are available at https://www.bioinfotiget.it/gitlab/custom/fm-benchmark. The fmaudit package (input-scale, fine-tuning, train/test-overlap and target-detection checks, task baselines, the expression-matched specificity test and a scorecard) is installable from the same repository and includes tests and a tutorial. The within-cell-type permutation null is provided as an analysis script in the same repository. The tables underlying each figure are provided as Supplementary Tables.

## Author Biographies

Francesca Cupaioli is a researcher at CNR-ITB working on single-cell transcriptomics and models of neurological disease.

Ivan Merelli is a senior researcher at CNR-ITB working on bioinformatics methods for genomics and single-cell analysis, high-performance computing and machine learning in biomedicine.

