## Supplementary_Methods for "Where single-cell foundation models help: a public-data benchmark of annotation, in silico perturbation and gene regulatory inference"

This file gives the full protocols summarised in the main-text Methods. Reference numbers refer to the main-text reference list. Supplementary Tables are in the Supplementary Tables file, and code, configurations and SLURM scripts are in the fm-benchmark repository (methods supplement in the repository documentation gives every deviation and job identifier).

Benchmark design

The benchmark is a grid of tasks (annotation, perturbation, GRN) by datasets grouped into three levels (Figure 1). Methods were four FM families used zero-shot (scGPT whole-human checkpoint [1], Geneformer V1-10M [2], Nicheformer [3], scFoundation [4]) and, where applicable, after fine-tuning. We also ran the current Geneformer release (V2-316M, with 4,096-token input and its own gene dictionaries) with exactly the V1 protocols of each grid row, CellOracle [18] with a promoter base GRN, optionally augmented with STRING v12 physical interactions (confidence ≥ 700) [19], the supervised perturbation models GEARS [20], CPA [21] and scGPT fine-tuned for perturbation, the virtual-cell model STATE [22] (SE-600M cell embedding with the State Transition model trained on Replogle and Nadig CRISPRi screens, using the public checkpoint in which K562 is held out as a whole cell context, with the variant that saw K562 reported only as a contaminated upper bound), and non-FM baselines chosen per task. All methods in a grid row were run on the same cells, split, gene universe and metric. Random seeds were fixed. Configurations, hyper-parameters and SLURM scripts are in the repository.

#### Datasets

Level 1 (public, primary cells and cell lines): the Immune Human atlas (33,506 cells, 16 types, 10 batches) [23]. The Tabula Muris Senis lung (10x, 24,540 cells, 30 types) [24], with classes named by the authors' free annotation. Norman K562 CRISPRa Perturb-seq (GEARS processing, 91,205 cells, 5,045 genes) [11] and Adamson K562 CRISPRi Perturb-seq (68,603 cells, 5,060 genes, 87 conditions) [12]. The CRISPRi essential-gene Perturb-seq screens of K562 (310,385 cells, 8,563 genes, 2,057 perturbations, 10,691 control cells, 48 libraries) and RPE1 (247,914 cells, 8,749 genes, 2,393 perturbations, 11,485 control cells, 56 libraries) [13] as raw counts from the scPerturb collection [25], with the GEARS-processed versions of the same screens (log-normalised, 5,000 genes) for the supervised models. The genome-scale CRISPRi Perturb-seq of primary human CD4 T cells [14] (four donors, at rest and after 8 and 48 hours of stimulation, 11,526 targets in the authors' differential expression table), of which the authors' differential expression statistics and pseudobulk counts were used for all conditions and cells were extracted from the four resting cell-level files only (1,500 non-targeting cells per donor and up to 50 cells per selected target and donor, 113,720 cells). Level 2 (neural tissue): the public CHOOSE mosaic organoid CRISPR screen [15] (28,344 cells, 36 perturbed genes, 991 non-targeting cells, 16 cell types annotated by the authors), with the HNOCA atlas [26] as annotation reference, the CRISPRi (32,300 cells, 184 targets, 437 control cells, 4 libraries) and CRISPRa (21,193 cells, 100 targets, 434 control cells, 2 libraries) screens of iPSC-derived neurons [17] as raw counts from the scPerturb collection, and the in vivo Perturb-seq of 35 autism and neurodevelopmental risk genes in the developing mouse cortex [16] (GEO GSE157977, frameshift mutations delivered in utero at E12.5, cells profiled at P7, 18 pooled libraries). Level 3 (human tissue): EGFR-mutant non-small-cell lung cancer biopsies profiled by Smart-seq2 [28], with the HLCA core [29] as human lung reference.

#### Annotation

Cell embeddings were computed zero-shot for each FM from raw counts. Within a dataset, labels were transferred by k-nearest neighbours (k = 30, cosine) from a stratified 50% reference split to the query half (Immune, Tabula Muris), and, for the CHOOSE organoid screen, within the dataset (stratified 50/50 split, 14,172 reference and 14,172 query cells, Supplementary Table S18) and from a 50,029-cell HNOCA reference subsample (k = 30,

Euclidean, evaluated as broad-lineage concordance with the authors' annotation through a crosswalk fixed in advance, with acceptance of the HNOCA label Glioblast for radial glia and astrocytes as a sensitivity analysis). Across datasets (HLCA core to the lung-cancer biopsies) and across species (Tabula Muris lung to the biopsies, mouse to human, HLCA to Tabula Muris lung, human to mouse), every model embedded reference and query from raw counts, labels were transferred with k = 20 and Euclidean distance, query cells with a mean neighbour distance above the 90th percentile and a vote fraction below 0.5 were left unassigned (the override), and accuracy was computed on an explicit crosswalk of shared coarse classes. Cells left unassigned, or predicted as a reference class without counterpart in the query, were not scored. Supplementary Table S3 gives their fraction and the accuracy obtained when they are counted as errors (for Nicheformer from human to mouse, 4.2% and 0.927 with the primary protocol, 0.1% and 0.978 with cosine kNN). Mouse genes were mapped to human one-to-one orthologs (17,105 pairs). Baselines were PCA (50 components of 2,000 highly variable genes) with the same kNN, and CellTypist [30] trained on the same reference. We report accuracy and macro-F1 with 95% bootstrap confidence intervals over query cells. As a control for the classifier, both cross-species directions were repeated with cosine kNN (k = 30, no distance or confidence override) for every embedding and for three baselines: PCA with global or per-dataset scaling and per-dataset z-scored highly variable genes (Supplementary Table S12). From human to mouse the primary protocol left 4.2% of Nicheformer's query cells unscored (0.927 counting them as errors), whereas cosine kNN scored 99.9% and reached 0.979 (macro-F1 0.919). Tabula Muris classes were named with the authors' free annotation, because the CELLxGENE field labels 2,424 endothelial cells as smooth muscle. The species represented in each FM's pre-training corpus were taken from the original publications and model cards (Supplementary Table S9).

Nicheformer was applied with the tokenisation of its repository tutorial for dissociated data: raw counts on the 20,310 model genes were scaled to 10,000 counts per cell, divided by the dissociated technology mean, ranked and prefixed with species, assay and modality tokens. Species was set per dataset (human, or mouse for Tabula Muris), modality to dissociated, and assay to the 10x chemistry of each library (3' v3 for CHOOSE, 3' v2 with the mouse species token for the mouse cortex, 3' v2 for Tabula Muris and Norman, 3' v1 for Adamson, per cell for HLCA and the immune atlas, and Smart-seq v4 as the nearest token for the Smart-seq2 biopsies). An earlier version of our pipeline omitted the species and modality tokens and used normalised values. Repeating every Nicheformer analysis with the tutorial settings changed its embeddings (median cosine similarity 0.84) but not the conclusions (Supplementary Table S91). Its K562 specificity runs and its GRN were not rerun and keep the earlier values.

The scGPT results of the main grid used the continual-pretrained whole-human checkpoint released for zero-shot cell embedding. Organoid screen annotation and knockouts were repeated with the scGPT brain checkpoint and with the original whole-human checkpoint, with identical scripts, cells, inputs and scoring (Supplementary Tables S18 and S92).

Fine-tuned annotation. For Geneformer V1, Geneformer V2, Nicheformer, scFoundation and scGPT, cell-type classifiers were fine-tuned on every annotation reference with a linear probe on frozen embeddings, partial unfreezing and LoRA, and for scGPT also the full fine-tuning protocol of its annotation tutorial. Each classifier was trained on a stratified subsample of 5,000 reference cells, of which 1,000 were held out for validation, with a reduced grid (two partial-unfreezing and two LoRA configurations per model and task, and the complete linear-probe grid), and the configuration with the highest validation macro-F1 was applied once to the test cells of the zero-shot analysis with the same coarse crosswalks. Logistic regression on 50 principal components, logistic regression on 2,000 z-scored variable genes and CellTypist, with its regularisation selected on the same validation cells, were trained on the same cells. Differences were assessed on the same test cells with a paired bootstrap of macro-F1 and an exact McNemar test, with Benjamini-Hochberg correction (Supplementary Figure S3, Supplementary Table S17). Because the fine-tuned classifiers and these baselines used 4,000 training cells, a 5,000-cell zero-shot kNN is also reported as a control for reference size. For the CHOOSE screen the non-targeting cells were excluded from the fine-tuning pool, PCA-kNN was added to the same-split baselines, the test cells were the 14,172 query cells of the zero-shot split, and only the continual-pretrained scGPT checkpoint was fine-tuned. Both choices were fixed before scoring (Supplementary Tables S7 and S19).

### In silico perturbation

Zero-shot FM perturbation followed the embed, delete and re-embed protocol: each target gene was removed from (or zeroed in) the input of each cell, and the effect was the cosine distance between original and perturbed cell embeddings, averaged per cell type. Before simulation, fmaudit's target-detection check reports, per target and cell type, the number and fraction of expressing cells and flags cell types below 10 expressing cells or 1% (defaults). scGPT samples 1,199 genes when a cell expresses more and breaks ties of its 51-bin quantile binning at random. Both steps were fixed per cell (seed = cell position), the input was the target (if expressed) plus the first other expressed genes of the fixed permutation, and the perturbed input was the same set without the target, with bins recomputed. Cells not expressing the target have identical inputs and zero effect. CellOracle simulated knockouts by signal propagation (three steps) on cluster-specific ridge GRNs and reported the per-cell-type norm of the simulated shift. In K562 the same protocol was applied to 500 control cells per dataset for every single-gene perturbation in the model vocabulary. Ground-truth effect size was the number of differentially expressed genes and the energy distance of perturbed vs batch-matched control cells, and direction agreement was the cosine between predicted and real embedding shifts. Because the protocol only models loss of function, a model capturing the correct axis should give negative cosines for CRISPRa and positive ones for CRISPRi. Supervised models were trained on the GEARS simulation split (seed 1) and scored by Pearson correlation of predicted and observed expression changes (Pearson δ) on unseen test perturbations. Baselines were the mean training-perturbation shift and, for combinations, the additive sum of single effects [7].

CHOOSE organoid screen. The authors' processed object was used. Knockouts of the 36 genes were simulated in all 991 non-targeting cells, for seven zero-shot models and checkpoints and eleven fine-tuned backbones, with the adapters of the annotation track (Geneformer rank tokeniser of the fine-tuning code, Nicheformer genes converted to Ensembl identifiers through the Geneformer dictionary, batched scFoundation forward and deterministic 1,200-gene scGPT input, Supplementary Tables S33 and S35). Ground truth per gene and cell type was the number of differentially expressed genes at BH FDR $< 0.05$ (367 testable pairs) and the absolute library-stratified Cochran-Mantel-Haenszel log odds ratio of composition (498 pairs). Predictions were scored by Spearman ρ pooled over pairs with bootstrap intervals, against naive predictors (target mean expression and detection rate, median library size and genes detected, number of control cells per cell type), a null that permutes perturbed genes within expression decile and a null that permutes them within cell type (1,000 permutations each), with the decision rule of the analysis plan ($\rho > 0$ with BH $q < 0.05$ in the family, at least 0.10 above the best naive predictor with a bootstrap interval of the difference above 0, and permutation $p < 0.05$). The previous per-gene statistic (median ρ over cell types with a sign-aware Fisher combination) is reported next to it. The replication ceiling was the agreement between ground truths computed from two halves of the libraries (four libraries per half), with the Spearman-Brown reliability of the full data (Supplementary Table S4). Cosine distances were computed in float64. Lung-cancer biopsies [28]: 23,261 Smart-seq2 cells. Ground truth was the number of differentially expressed genes per cell type for residual disease versus treatment-naive biopsies, computed per cell (Wilcoxon) and, as a patient-aware control, by pseudobulk edgeR. Progression versus treatment-naive served as a specificity control. Library size and number of genes detected per cell type served as complexity baselines.

In vivo Perturb-seq of the mouse cortex. The GEO deposit (one archive, 18 pooled expression libraries, 18 dial-out libraries and two single-perturbation libraries, Cell Ranger matrices on mm10 with raw UMI counts) holds neither the barcode-to-gene table nor cell types, so the authors' per-cell annotations (cell class, perturbation, batch, key-cell flag) were taken from the public viewer of Single Cell Portal study SCP1184 [27] and matched to the GEO matrices by library and cell barcode (49,067 of 49,067 cells). Authors' batches were matched to GEO libraries by barcode overlap. An independent reconstruction from GEO alone agreed with the authors' classes for 97.7% of key cells and with their perturbation calls for 99.3% of analysis cells. Analysis cells were the key cells with one of the 35 genes or the GFP control (Supplementary Table S2, 17,889 cells, 17 libraries, five classes: excitatory and inhibitory neurons, astroglia, microglia and oligodendrocyte lineage). The GFP control has 374 cells. Ground truth: raw counts summed per class, library and perturbation, edgeR quasi-likelihood [32] with a paired design (library plus group), filterByExpr and TMM, number of genes at FDR < 0.05 and summed absolute log2 fold change over tested genes. The planned reference was GFP cells of the same class and library (a library enters with at least 3 perturbed and 3 GFP cells). The strict scorable rule of the plan (at least 20 perturbed cells in each of 2 libraries) gave 7 pairs, below the threshold written in advance, so the pooled reading (at least 20 perturbed cells summed over at least 2 libraries) was primary (72 pairs, 32 genes, 4 classes). A secondary reference was all other analysis cells of the same class and library (152 pairs, Supplementary Table S5). Composition was tested by Mantel-Haenszel log odds ratio across libraries. The replication ceiling was computed from odd and even GEO libraries. Knockouts used all 374 GFP cells from raw counts (sensitivity: 1,170 cells of the GFP-only library). Mouse genes were mapped to human one-to-one orthologs with the table and code of the cross-species annotation (32 of the 35 targets present, Pten, Stard9 and Tcf20 absent). Nicheformer used the mouse, 10x 3' v2 and dissociated tokens. Old gene names were matched by alias (Mll1 to Kmt2a, Myst4 to Kat6b, Scn2a1 to Scn2a). The adapted knockout scripts reproduced stored CHOOSE per-cell values (maximum absolute difference 2.4e-07, Supplementary Table S93). Scoring followed the organoid screen, with the null permuting genes within cell class (primary family: five models by two truths). CellOracle and the scGPT brain checkpoint were not run in vivo. On the 14 genes shared with CHOOSE, the measured burden of the two systems was compared, and the within-cell-type null was also applied to the stored CHOOSE predictions (Supplementary Tables S40, S43 and S44).

Geneformer native protocol. Geneformer was also run with its package in silico perturbation protocol: raw-count rank-value inputs from the package tokenizer, second-to-last-layer cell embeddings (mean-pooled for V1, the classification token for V2), overexpression by moving the target to the first rank for CRISPRa, and the goal-state shift, the change in cosine similarity between a cell and a target-state centroid after the perturbation. In Norman, each of the 152 single targets and 50 detection-matched control genes was overexpressed in 500 control cells.

Supervised perturbation fine-tuning. CHOOSE was converted to the GEARS format on log-normalised expression in the 5,036-gene space of the CHOOSE analysis, and the 36 genes were split into five gene-level folds before training. GEARS and scGPT fine-tuned for perturbation prediction were trained with the settings of the K562 screens and scored by Pearson δ on all genes and on the 20 most differentially expressed genes, pooled and per cell type, against a training-mean baseline, a no-change baseline and the naive expression baselines (one-sided paired Wilcoxon tests over genes, Supplementary Table S52). scFoundation was combined with GEARS as in its original publication, with the authors' code, checkpoint, hyper-parameters and preprocessed Norman and Adamson data, and frozen scFoundation gene-context embeddings as GEARS input. The GEARS test split was mapped onto these data by perturbation name (99 of the 106 unique Norman and 20 of the 22 Adamson test perturbations), and the same code was run without the embedding as a matched comparator (Supplementary Table S31). On CHOOSE it was trained on the same five folds. Genes outside its 19,264-gene list were given the observed control value, and the 3,056 shared genes are reported as a sensitivity analysis. Geneformer V1 and V2 were fine-tuned as perturbation classifiers on the training knockouts, the only fine-tuning route provided by its authors, and then used for in silico deletion of the held-out genes, with a classifier trained on permuted labels as negative control. V2 was trained for one epoch instead of ten, for compute reasons (Supplementary Table S53). Nicheformer provides no perturbation or gene-embedding interface and was not fine-tuned for perturbation.

Genome-scale screens. T cells: eligible targets were present in the authors' differential expression table in all three conditions, with significant on-target knockdown and without off-target flags in all three, and scored in at least three donors in every condition (5,217 of 11,526). Sixty targets were drawn per decile of target log CPM in the resting non-targeting pseudobulks (numpy RandomState(0), candidates sorted by Ensembl identifier, Supplementary Tables S94 and S95). The 200 control cells were the first 50 non-targeting cells with a single guide of each donor in a seeded permutation. Every model used raw UMI counts (median 7,650 UMI and 3,560 genes per control cell, Supplementary Table S96), with the model settings of the K562 analyses: deterministic input for scGPT, the package tokenisers of Geneformer V1 (2,048 tokens) and V2 (4,096 tokens), the tutorial tokenisation of Nicheformer (assay token 10x transcription profiling, because the probe-based kit has no token of its own) and the 19,264 genes of scFoundation. Per-donor effects were recomputed with edgeR (TMM, dispersions estimated on the non-targeting pseudobulks, likelihood-ratio test of the guides of a target against all non-targeting guides, FDR 10%, Supplementary Table S97). Direction was the cosine between the predicted shift and the shift of up to 50 real perturbed cells, compared with the shifts of 100 other targets. Specificity used 20 control genes per target from a pool of 1,200 genes matched for expression. Supervised models used up to 100 perturbed cells per target and 2,000 control cells (5,000 highly variable genes plus the targets) with five folds over the 600 targets, and networks were extracted from 2,000 non-targeting resting cells for hubs among the regulators of the CellOracle promoter base GRN with at least 50 DEGs, scored against their DEGs and, as a secondary truth, against DEGs replicated in at least three donors. Cell lines: real effects were computed from log1p counts per 10,000 on the 3,000 genes of highest combined variance, with up to 3,000 control cells sampled per library in the proportions of the perturbed cells (seed 0). Eligible essential-gene targets were present in both K562 and RPE1 with at least 25 perturbed cells and measured in both (1,550 of 2,055 shared), with 60 drawn per decile of K562 control expression. Neuron screens used all targets with at least 10 perturbed cells. Up to 100 real perturbed cells per target entered the direction endpoint. For Geneformer V2 and scFoundation, which cost more than 30 ms per cell, K562, RPE1 and neuron CRISPRa were run on 100 control cells and 20 real cells per target, a rule fixed from measured cost before scoring (Supplementary Table S98). On the models run with the full set, effects on the 100-cell subset correlated with the full effects at ρ 0.94 to 0.99. Geneformer V2 used bf16 in these runs after a precision check against fp32 ($\rho \geq 0.999$). Supervised models used the GEARS simulation split (seed 1, train gene fraction 0.75) on the perturbations whose target is among the 5,000 genes (K562 278, 31 and 103, RPE1 439, 49 and 163 training, validation and test perturbations), with GEARS (hidden size 64, 20 epochs) and scGPT with the GEARS pipeline (whole-human checkpoint, 15 epochs), and were scored by Pearson δ on the top 20 DE genes against the training mean and a split-half ceiling. scFoundation with GEARS was not run on these screens because it needs the 19,264-gene input rebuilt from raw counts and more than a day of GPU per screen.

Gene representations and zero-shot re-read. Context-free gene vectors were read from the checkpoints used throughout: the scGPT encoder gene embedding before its normalisation, the Geneformer V1 and V2 word embeddings, the Nicheformer input token embedding and the scFoundation gene embedding, without positional embeddings, matched to targets by Ensembl identifier or symbol. Transfer splits used two halves of the shared targets fixed by the MD5 hash of the Ensembl identifier, training on one half with the truth of the training system and testing on the other with the truth of the test system, then the reverse, and the primary family (5 FMs by 4 splits) used the decision rule of the plan with random normal vectors of the same dimension (5 seeds) as an additional control. For the confirmatory genome-wide screen, a copy of the raw counts was checked against the original file, effects were computed for all 7,568 eligible targets as in the essential-gene screens, and the folds were fixed by the MD5 hash of the Ensembl identifier. The first scoring script was run on Poisson noise truths as a code test, which showed gains of FM plus expression over expression of up to 0.05 and a permutation of gene-vector rows that did not reproduce them, so the final scoring was cancelled before it started and the plan was amended to a Freedman and Lane null (200 truths) with a secondary null of 100 permutations of the truth. The same null was applied to the earlier within-system results. The nonlinear baseline (each expression feature and STRING degree as linear, squared and decile-indicator terms) was added after the primary results and is reported as a robustness analysis. For the zero-shot re-read, control cells not expressing the target were verified to have exactly zero change in every model, targets with fewer than five expressing control cells were excluded, and in Geneformer V1 and Nicheformer the target is absent from part of the expressing cells because of input truncation, so their effect per expressing cell still contains zero distances by construction. The genome-wide scoring was repeated for Geneformer V1-10M, Geneformer V2-316M and scFoundation with only the feature matrix changed, targets without a vector dropped per model (11, 10 and 72 of 7,568) and the expression baseline refitted on the same targets, with BH over these three tests and the same null. In the smaller essential-gene and T cell sets the Geneformer gains were at most 0.024, with intervals that included 0, and scFoundation gains were negative. Direction in the genome-scale screens. For each target the predicted shift was the mean deleted minus original embedding over the stored control cells, and the real shift the centroid of up to 50 perturbed cells minus the batch-matched control centroid, embedded with the same model and input pipeline. The statistic D was the median over targets of the cosine between predicted and real shift minus the mean cosine of the predicted shift with the real shifts of 100 other targets, tested by a one-sided sign-flip permutation (10,000 flips) with BH over 20 tests (five models by four screens). On 100 targets per test, drawn once with seed 0, the same cosine was compared with the mean cosine of the shifts predicted for 20 control genes matched for detection-rate decile and mean expression and deleted in the same cells (paired bootstrap), and a test was positive when both comparisons were. The reused deletion scripts reproduced the stored predicted shifts on 10 targets per model and screen, the earlier secondary direction scores are reported unchanged, and the direction of partial knockdown could not be assessed because those runs stored no shift vectors (Supplementary Table S89). Partial knockdown. For each target and screen the residual expression r was the ratio of its normalised expression in perturbed and matched control cells (in T cells, 2 to the authors' log2 fold change at rest), clipped to the interval from 0 to 1 (median 0.14 in T cells, 0.17 in K562, 0.09 in RPE1 and 0.72 in neuron CRISPRi). The target count was scaled by r in the control cells before each model's own input step: re-binning for scGPT, re-ranking with the package tokenizer for Geneformer, re-tokenising for Nicheformer and re-normalising for scFoundation. All other inputs, checkpoints, control cells and distances were those of the stored deletion runs, which the modified scripts reproduced on 20 targets per run, and with r = 1 no input changed. The knockdown effects were scored with the original decision rule, a paired bootstrap against deletion, the Freedman and Lane incremental-value test and, as a sensitivity analysis, a fixed r of 0.2 (Supplementary Table S99). Against deletion on the same targets (paired bootstrap), knockdown changed ρ in 4 of 15 cell-line tests, lower for scGPT (−0.117) and scFoundation (−0.120) in K562 and higher for Geneformer V1 in K562 (+0.013) and RPE1 (+0.012), and in none of the T cell tests. Knockdown effects of scGPT and scFoundation differed strongly in size from deletion effects, because a scaled count is a new input value for these encodings, whereas Geneformer V1 and Nicheformer ranked targets nearly as with deletion.

Newer models. Up to four models with public weights and a peer-reviewed publication from 2024 to 2026 were ranked, before any score, by the number of independent published benchmarks that include them (web survey of seven benchmark studies, 22 November 2026): UCE 33-layer, SCimilarity v1.1, CellPLM 85M and TranscriptFormer Exemplar, each with the authors' checkpoint and preprocessing, run on an L40 GPU (SCimilarity on CPU). UCE, SCimilarity and CellPLM had released weights in 2023 as preprints, and with first release of weights as the criterion CellFM, scPRINT and AIDO.Cell would have replaced them. They were scored with the unchanged cross-species and within-dataset annotation protocols (FM accuracy above the best baseline under the same protocol, paired bootstrap of query cells) and the confirmed gene-vector protocol, with BH over the 19 primary tests, and with the cell-line deletion rule on the 600 K562 essential targets as a secondary analysis. SCimilarity has no gene-level output and UCE samples its input genes at random, so SCimilarity was not scored for gene vectors and UCE not for deletion. UCE covered 7,422 of 7,568 genome-wide targets. In the human-to-mouse direction the plan compared each model with the best baseline of the original protocol (CellTypist with default settings), and the tuned CellTypist value is reported alongside. TranscriptFormer reached 0.264 from mouse to human with Euclidean neighbours but 0.934 with cosine neighbours. Recomputed kNN baselines equalled the earlier values exactly (Supplementary Table S90).

### Gene regulatory networks

For every FM we derived two gene-gene networks per system from the same 3,000 cells: cosine similarity of contextual gene embeddings and symmetrised attention averaged over layers and heads. We also evaluated the attention of scGPT fine-tuned on Adamson, CellOracle links (coefficient magnitude) and three baselines: Pearson co-expression, an expression prior (ranking genes by mean expression) and random ranking. For each hub gene, the top-50 neighbours were scored against a truth set within a common gene universe: in K562, genes differentially expressed after knockdown of the hub (Adamson) and ChIP-seq targets in K562 from ChIP-Atlas [34]. In the lung-cancer biopsies, genes responding to EGFR inhibition in treatment-naive cancer cells. The metric was fold enrichment over expectation after matching for expression level (FE@50, 1 = chance), pooled over hubs.

Networks from fine-tuned backbones were derived with the same protocol from scGPT fine-tuned for perturbation on Adamson or Norman, and from each of the five models fine-tuned on K562 cells to classify 76 Adamson perturbations and control cells. The ten K562 hub perturbations were excluded from this fine-tuning, and its training cells were disjoint from the network cells and from the control cells that define knockdown targets. Networks were scored with the unchanged expression-matched FE@50 and compared with the zero-shot network of the same model and with co-expression by a bootstrap over hubs (Supplementary Table S51).

### Fine-tuning

Knockouts with fine-tuned backbones. In the organoid screen, knockouts were recomputed with the partially unfrozen and LoRA backbones of every model selected for cell-type classification, and for scGPT also with its tutorial full fine-tuning (11 backbones). Linear-probe backbones are identical to the pretrained ones and were not rerun. Weight changes were verified tensor by tensor in every arm (expected tensors changed, frozen tensors bit-identical). Fine-tuned scGPT backbones were evaluated without the classification decoder. The Geneformer V2 zero-shot table is fp32 and its fine-tuning adapter bf16. Effects were compared with zero-shot by median ratio and rank correlation over pairs, and their agreement with the ground truth by the change in $\rho$ with a bootstrap interval (Supplementary Tables S37 and S38).

### Expression-matched specificity test

To ask whether an in silico knockout carries information about the deleted gene beyond the removal of expression, we compared, in each cell, the embedding shift after deleting the target g (the observed effect, computed with each model's existing protocol) with the shifts after deleting, one at a time, $K = 20$ control genes matched to g in that cell (same expression rank window for Geneformer, same expression bin for scGPT, Nicheformer and scFoundation, and same detection-rate decile across the dataset, controls in the model vocabulary, excluding STRING partners of g and genes sharing a family prefix). The specific effect is $S = \text{observed} - \text{mean(control)}$, and $z = S / \text{sd(control)}$. Per-cell-type values average over cells expressing g, and pair-level $z > 2$ marks a gene×cell-type pair with a specific excess shift. For CellOracle, whose knockout acts on the whole network, the controls were 20 active regulators matched on detection and mean expression at dataset level. The organoid screen used its 991 control cells, with the matched controls of Geneformer V2 and scFoundation computed on at most 200 expressing cells per gene for compute reasons and the same cap applied to the other models as a post hoc sensitivity analysis. K562 used 200 control cells per screen. The fraction of pairs in which the target exceeds all 20 controls (4.8% expected by chance) was tested with a binomial test (Supplementary Tables S6 and S45).

### Statistics and computation

Correlations are Spearman's $\rho$ with two-sided p-values unless stated otherwise. Multiple testing was controlled with the Benjamini-Hochberg procedure [35] within pre-specified families: K562 zero-shot tests, organoid screen annotation (135 McNemar tests), organoid screen knockouts (36 tests of the decision rule, Supplementary Table S7) the mouse cortex (10 primary tests), the T cell screen (15 tests of five models by three conditions) and the cell-line screens (20 tests of five models by four screens). The analyses of the organoid screen, the mouse cortex and the genome-scale screens followed an analysis plan written before their results, and choices that the plan left open were fixed from the design and recorded before scoring. All computation ran on SLURM nodes (NVIDIA A100 and L40 GPUs) at CNR-ITB.

### Supplementary Tables

The Supplementary Tables are provided as one Excel file. Its Contents sheet lists every table with its title, description and size, each table is on its own sheet, and tables are numbered in order of first citation in the main text and the Supplementary Methods.

### Supplementary Figure legends

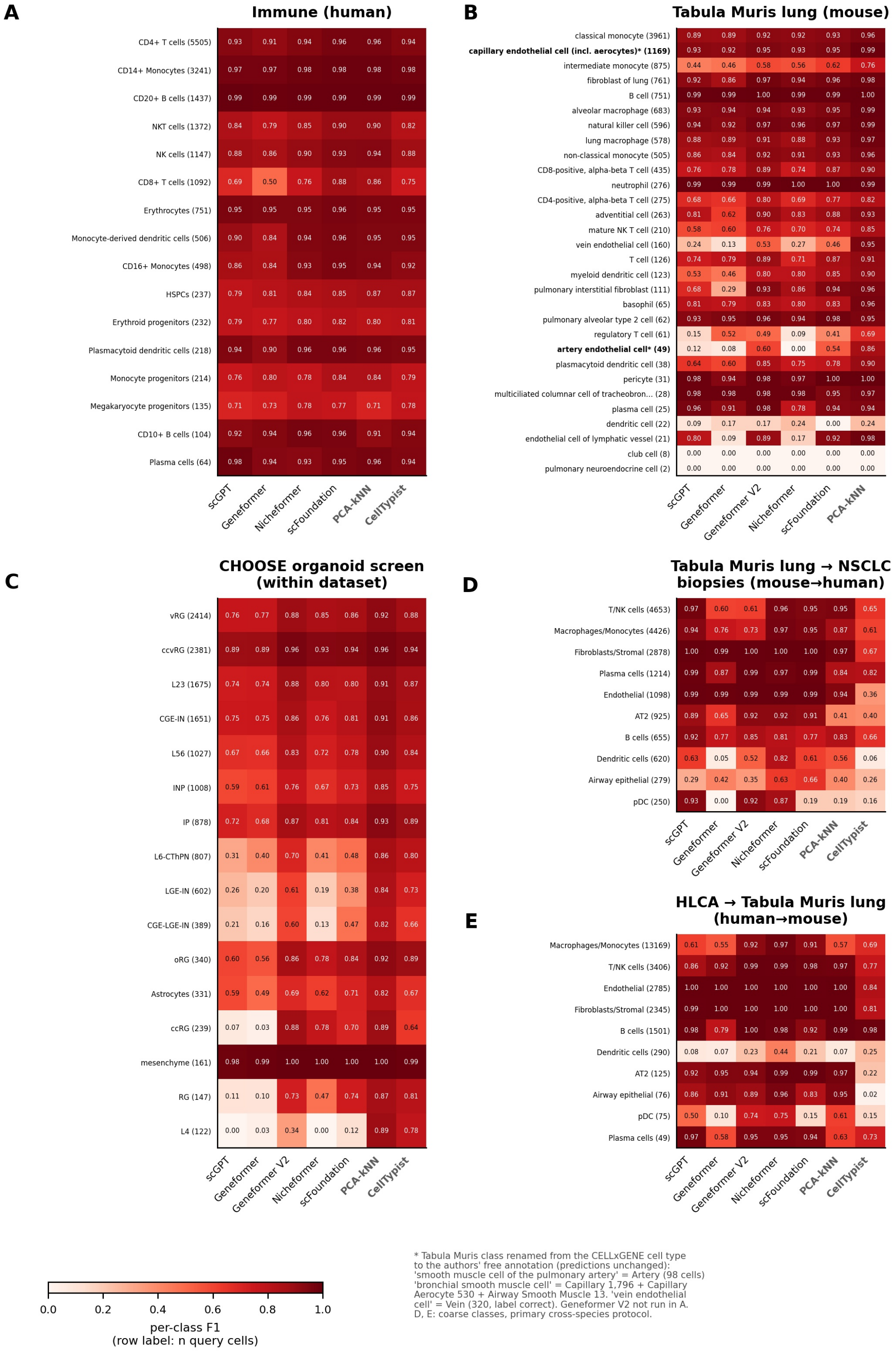


**Supplementary Figure S1.** Per-class F1 for all annotation systems. (A) Immune atlas. (B) Tabula Muris lung. (C) CHOOSE organoid screen within the dataset (16 author cell types, 50/50 split). (D) Tabula Muris lung to NSCLC biopsies (mouse to human). (E) HLCA core to Tabula Muris lung (human to mouse). Row labels give the number of query cells. D and E use coarse classes and the primary cross-species protocol. Geneformer V2 is shown where per-class predictions were computed. * Tabula Muris class renamed from the CELLxGENE cell type to the authors' free annotation, predictions unchanged.

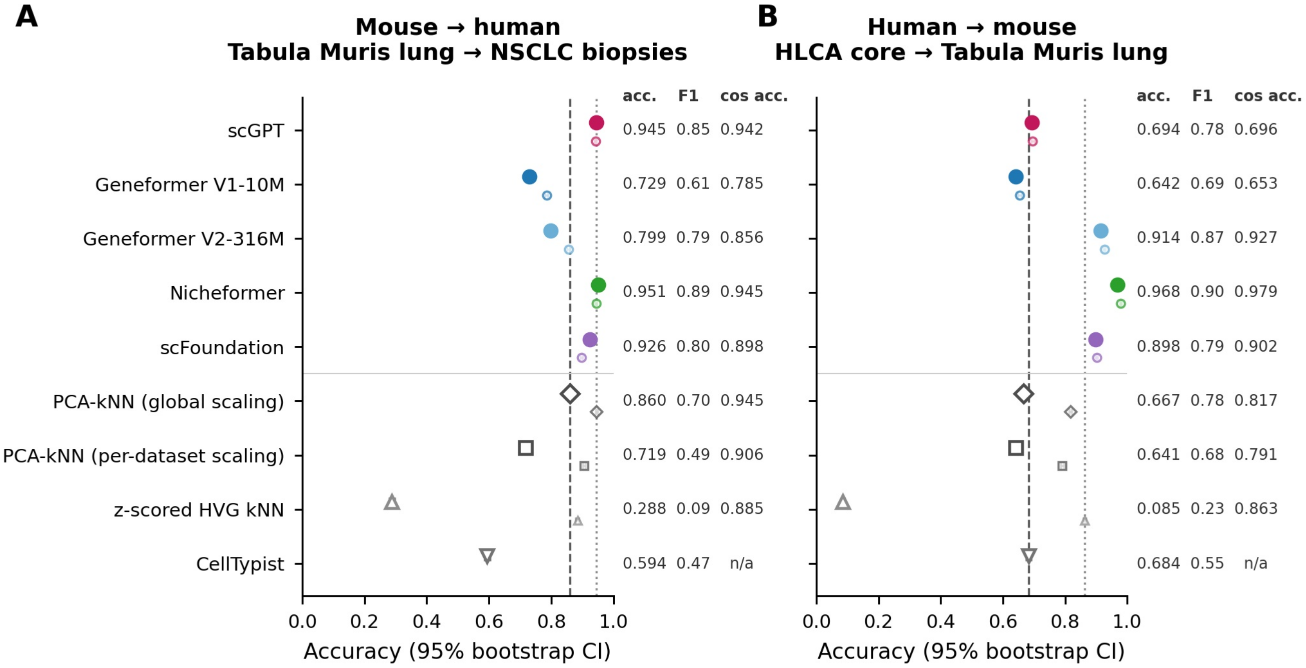


Large markers: primary protocol (kNN k = 20, Euclidean + override), filled = foundation models (zero-shot embeddings), open = non-FM baselines (ortholog gene space), dashed line = best non-FM baseline. Small markers: cosine kNN control (k = 30, no override), dotted line = best non-FM baseline under the cosine control. Coarse crosswalk (10 shared classes), CellTypist = custom logistic regression (no kNN). Labels: accuracy, macro-F1, cosine accuracy.

**Supplementary Figure S2.** Cross-species annotation with a cosine classifier control. Accuracy of kNN label transfer (A) from mouse Tabula Muris lung to human lung-cancer biopsies and (B) from human HLCA to mouse Tabula Muris lung, on shared coarse classes, for five FM embeddings and the non-FM baselines. Large markers, primary protocol (k = 20, Euclidean, with override). Small markers, cosine kNN control (k = 30, no override). Dashed and dotted lines, best baseline under each protocol.

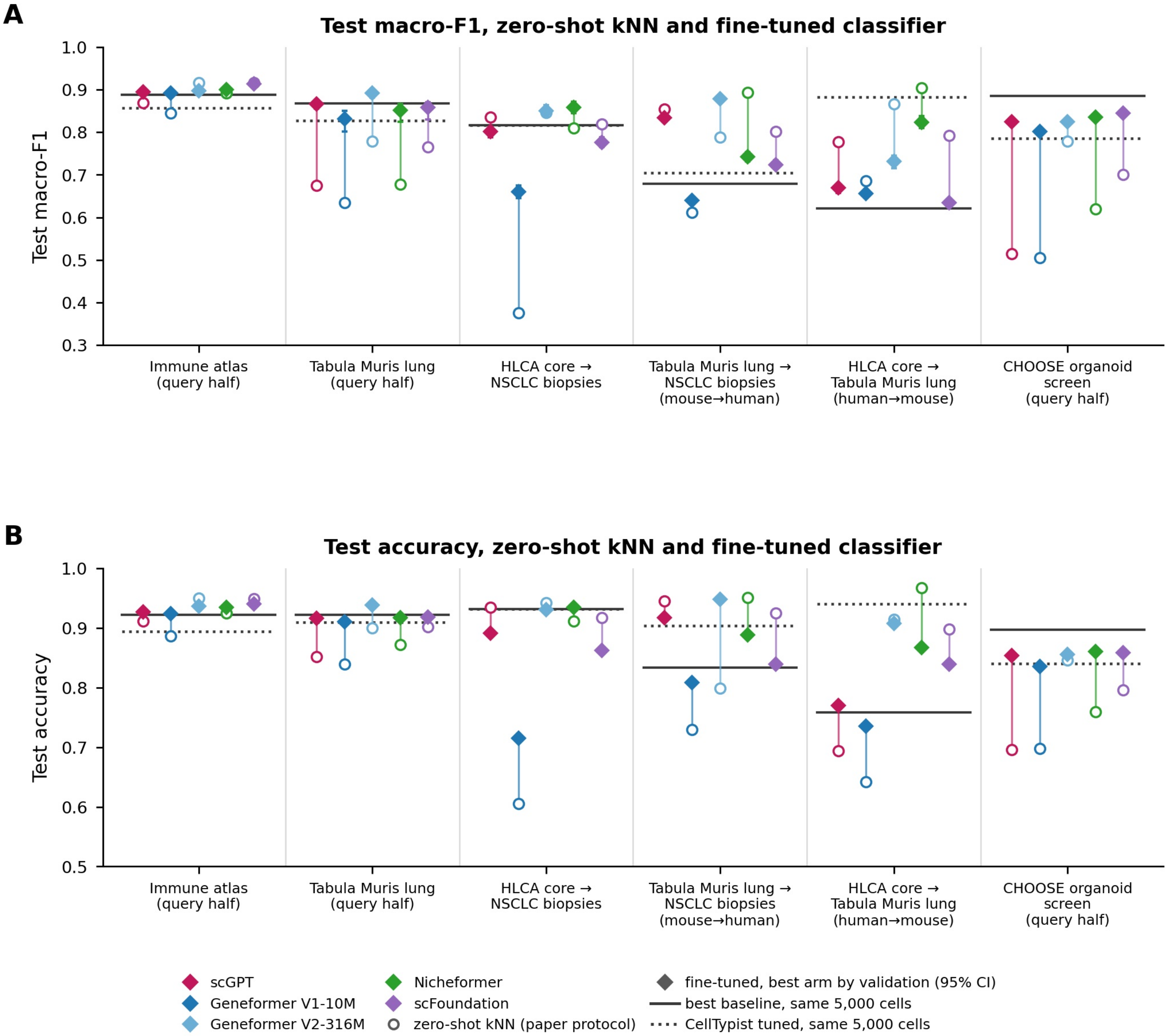


**Supplementary Figure S3.** Fine-tuned cell-type annotation on every annotation dataset. (A) Test macro-F1 and (B) test accuracy of zero-shot kNN on the full reference (open circles, values of Figure 2) and of the fine-tuned classifier of the same model (diamonds, 95% bootstrap interval) for the immune atlas and Tabula Muris lung query halves, HLCA core to NSCLC biopsies, the two cross-species transfers and the query half of the CHOOSE organoid screen. Classifiers used a linear probe, partial unfreezing or LoRA (scGPT also its tutorial full fine-tuning), chosen on 1,000 validation cells by macro-F1 and trained on 4,000 cells of a 5,000-cell stratified subsample of the reference. Solid line, best baseline trained on the same cells (PCA-LR, z-scored HVG-LR, CellTypist or, for the organoid screen, PCA-kNN, chosen on validation). Dotted line, CellTypist tuned on the same cells.

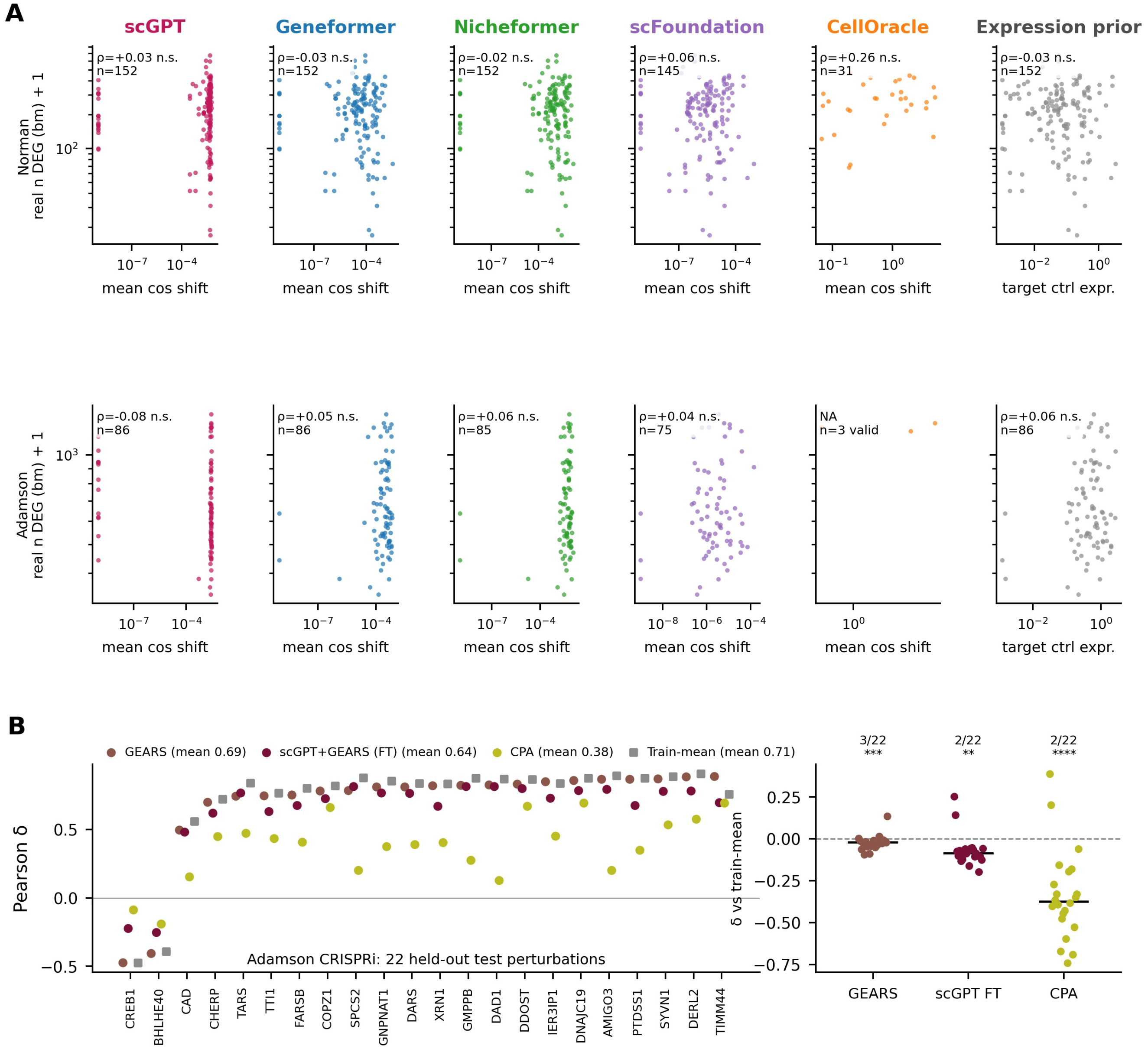


**Supplementary Figure S4.** K562 perturbations one by one. (A) Zero-shot predicted knockout magnitude (mean cosine shift) versus the real number of differentially expressed genes per perturbation, for each FM, CellOracle and the expression prior, in Norman (CRISPRa) and Adamson (CRISPRi). (B) Pearson δ of GEARS, scGPT+GEARS and CPA on the 22 held-out Adamson test perturbations, and their difference from the training-mean baseline.

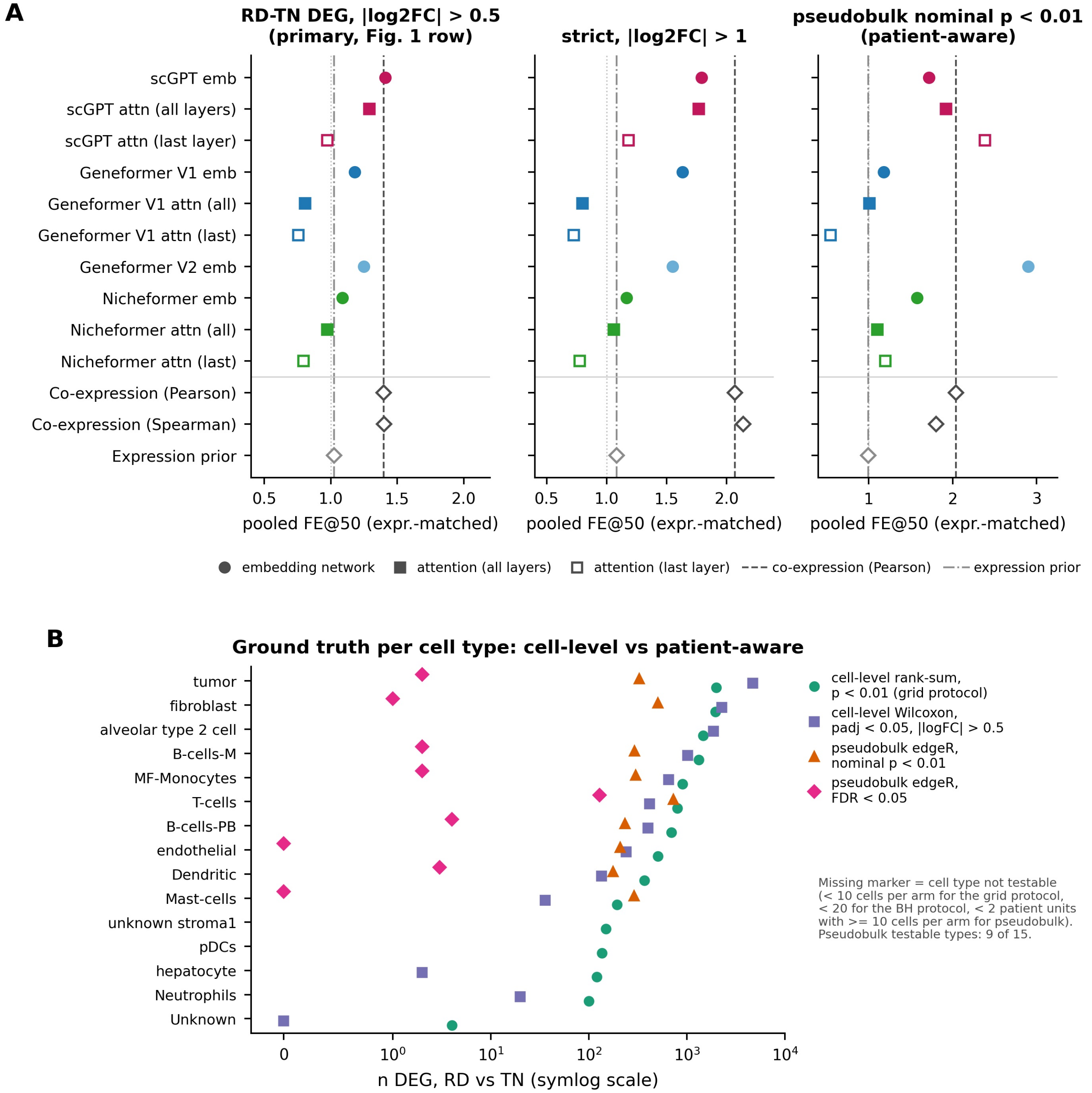


**Supplementary Figure S5.** EGFR-mutant lung-cancer biopsies. (A) Pooled expression-matched FE@50 of FM embedding and attention networks, co-expression and the expression prior against EGFR-TKI responsive genes under three definitions of the ground truth (cell-level, strict, patient-aware pseudobulk). (B) Number of differentially expressed genes per cell type, residual disease versus treatment-naive, computed at cell level and with patient-aware pseudobulk edgeR.

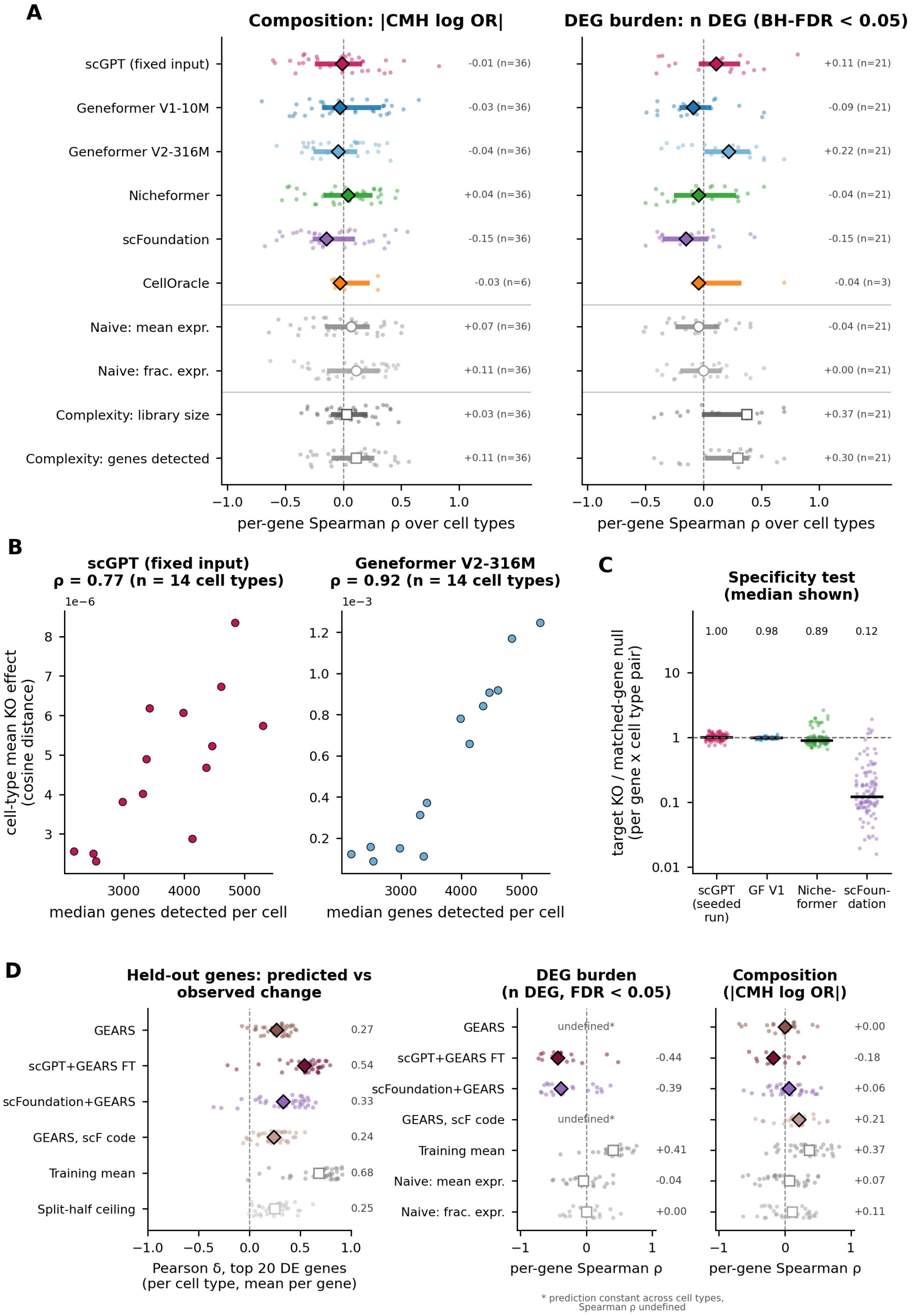


**Supplementary Figure S6.** CHOOSE mosaic organoid screen, earlier per-gene analysis. (A) Per-gene Spearman ρ over cell types between predicted knockout effect and composition change (|CMH log OR|) or FDR-corrected DEG burden, for FMs (scGPT with fixed input), CellOracle, naive expression predictors and cell-complexity baselines. (B) Mean predicted knockout effect per cell type versus median genes detected per cell for scGPT (fixed input) and Geneformer V2. (C) Expression-matched specificity test: ratio of target knockout to matched-gene control effect per gene×cell-type pair. (D) Supervised fine-tuning on the screen with five-fold gene hold-out: GEARS, scGPT with GEARS, scFoundation with GEARS and GEARS without embedding from the scFoundation code. Left, per-cell-type Pearson δ on the top 20 DE genes of each held-out gene, against the training-mean prediction and the split-half ceiling of the screen. Middle and right, the per-gene ρ of panel A computed from the fine-tuned predictions. Asterisk, prediction constant across cell types, so ρ is undefined. All fine-tuned models stay below the training mean, which carries the DEG burden per cell type shared by all knockouts.

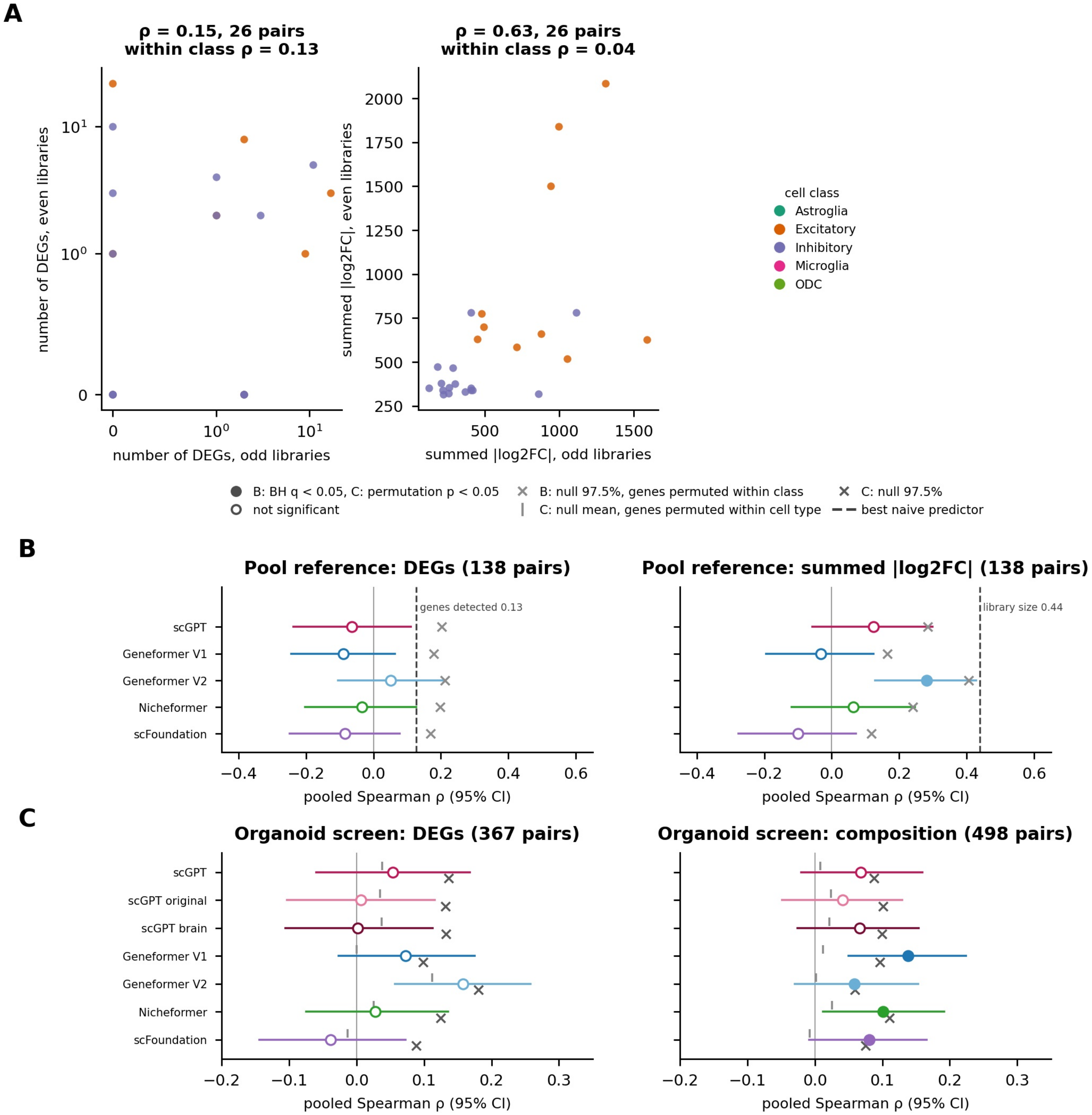


**Supplementary Figure S7.** Mouse cortex in vivo Perturb-seq. (A) Measured effect of each gene by class pair computed from odd against even GEO libraries (GFP reference, 26 pairs scorable in both halves): number of DEGs (left, symlog) and summed |log2FC| (right), with the Spearman ρ between halves and the ρ within cell class. (B) Secondary reference (all other perturbed cells of the same class and library, 138 scored pairs): pooled Spearman ρ (95% CI) of the five models with the number of DEGs and the summed |log2FC|, filled markers BH $q < 0.05$, crosses the 97.5% quantile of the null that permutes genes within class, dashed line the best naive predictor. (C) The same null applied to the stored predictions of the organoid screen (genes permuted within cell type): pooled ρ (95% CI), null mean (tick) and 97.5% quantile (cross), filled markers permutation $p < 0.05$.

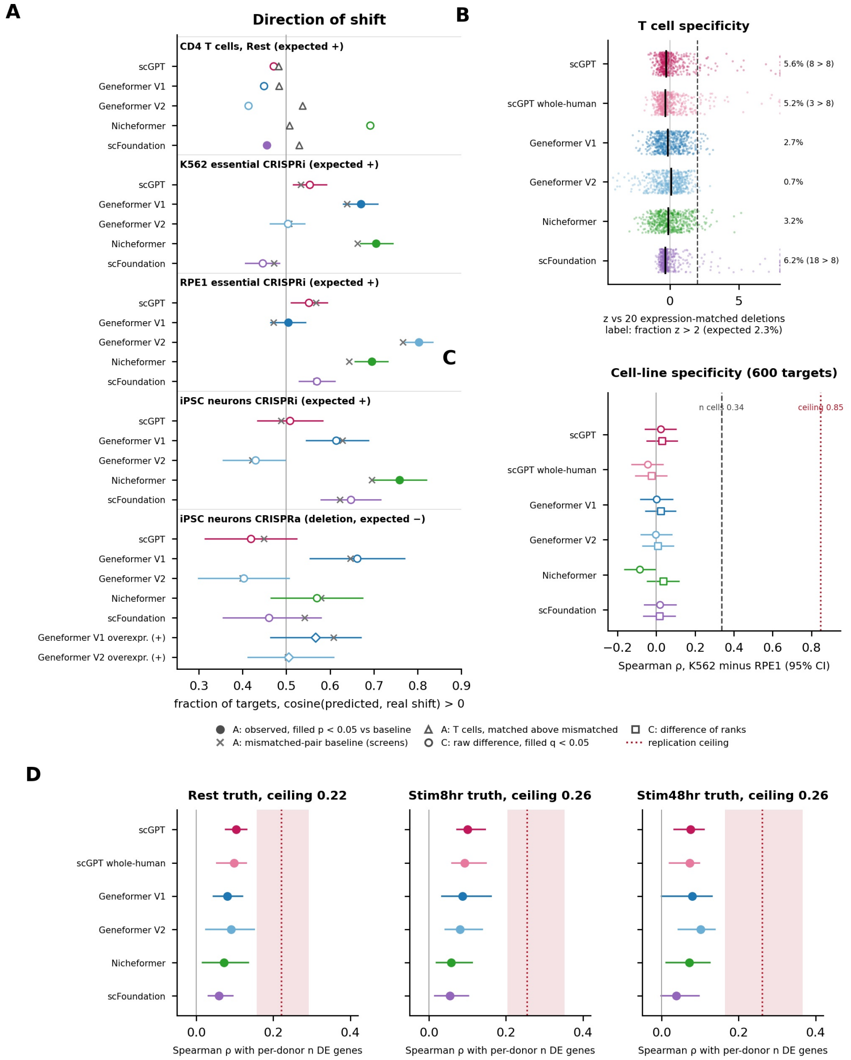


**Supplementary Figure S8.** Secondary endpoints of the genome-scale screens. (A) Direction. The fraction of targets whose predicted shift (mean embedding change over the control cells) has a positive cosine with the real shift (perturbed minus control centroid). For the scPerturb screens, the 95% bootstrap interval is shown and crosses mark the mismatched-pair baseline, which pairs each predicted shift with the real shift of another target. Filled markers mean the observed fraction exceeds that baseline at permutation $p < 0.05$. For T cells, triangles give the fraction of targets whose matched cosine exceeds the mean cosine with the real shifts of 100 other targets. Filled markers are Wilcoxon $p < 0.05$ for matched minus mismatched cosine. On the CRISPRa neuron screen the deletion is expected to point against the real shift, and the Geneformer overexpression rows are expected to point with it. (B) Expression-matched specificity in T cells: z of each target deletion against 20 expression-matched control-gene deletions in the resting control cells. Black bars are medians. Labels give the fraction of targets with $z > 2$ (2.3% expected) and the number of values above 8, which are drawn at the axis edge. (C) Cell-line specificity: Spearman ρ (95% CI) between the predicted K562 minus RPE1 effect and the real difference in the number of DE genes over the 600 shared targets. Circles are the raw difference and squares the difference of within-screen ranks. Filled markers would mean BH $q < 0.05$ (none). The dashed line is the best naive predictor (difference in the number of perturbed cells, ρ 0.34). The red dotted line is the replication ceiling (the real difference in library half A against half B, ρ 0.85). (D) Per-donor agreement in T cells: resting predictions against the per-donor edgeR number of DE genes in each culture condition. The dot is the mean over the 4 donors and the bar the range. The red dotted line and band are the mean and range of the six donor-versus-donor correlations (donor ceiling).

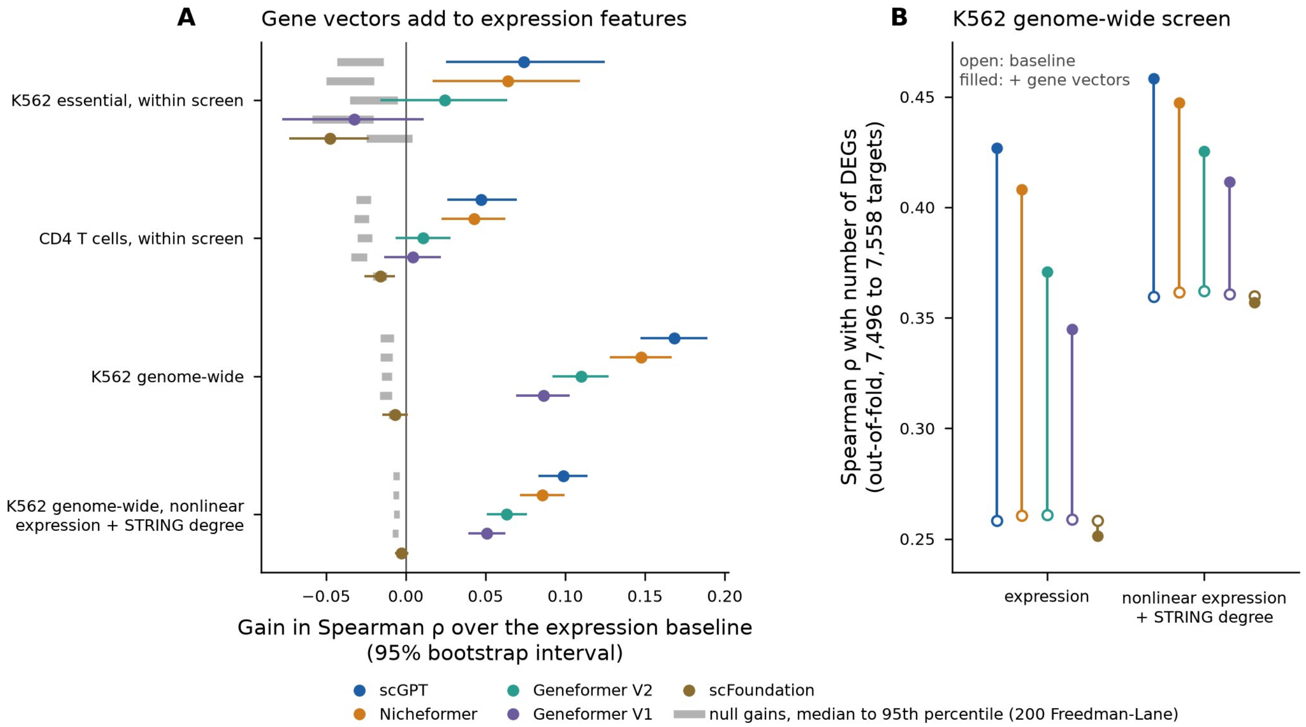
A
Gene vectors add to expression features
K562 essential, within screen
CD4 T cells, within screen
K562 genome-wide
K562 genome-wide, nonlinear expression + STRING degree
−0.05
0.00
0.05
0.10
0.15
0.20
Gain in Spearman ρ over the expression baseline (95% bootstrap interval)
B
K562 genome-wide screen
open: baseline
filled: + gene vectors
Spearman ρ with number of DEGs (out-of-fold, 7,496 to 7,558 targets)
0.45
0.40
0.35
0.30
0.25
expression
nonlinear expression + STRING degree
scGPT
Nicheformer
Geneformer V2
Geneformer V1
scFoundation
null gains, median to 95th percentile (200 Freedman-Lane)

**Supplementary Figure S9.** Gene vectors of foundation models predict knockdown effect size. (A) Gain in out-of-fold Spearman ρ with the number of DEGs when the context-free gene vectors of scGPT (blue), Nicheformer (orange), Geneformer V2 (teal), Geneformer V1 (violet) or scFoundation (brown) are added to expression features (target expression and detection in control cells, number of perturbed cells), in the K562 essential-gene screen and CD4 T cells at rest (within screen), the genome-wide K562 screen and, in the last row, against a nonlinear baseline that also includes STRING degree (added after the primary results). Points and bars, gain and 95% bootstrap interval over targets. Grey bars, median to 95th percentile of the gains of the whole procedure on 200 null truths with permuted residuals of the expression features (Freedman and Lane). (B) Genome-wide K562 screen (7,496 to 7,558 targets per model): out-of-fold ρ of the baseline (open) and of the baseline plus gene vectors (filled), for expression features and for the nonlinear baseline with STRING degree. Values in Supplementary Table S88.
